# Multiplexed ultrasound imaging using clustered gas vesicles and spectral imaging

**DOI:** 10.1101/2021.10.15.464596

**Authors:** Sangnam Kim, Siyuan Zhang, Sangpil Yoon

**Author notes:** Corresponding author: 151 Multidisciplinary Research Building, Notre Dame, IN 46556.

## Abstract

Current advances in ultrasound imaging techniques including super-resolution ultrasound imaging allows us to visualize microvasculature in biological specimens using microbubbles. However, microbubbles diffuse in blood stream limiting imaging acquisition and frame subtraction scheme of super-resolution ultrasound imaging cannot improve spatial resolution without moving microbubbles. Fluorescent proteins revolutionized to understand molecular and cellular functions in biological systems. Here, we devised a panel of gas vesicles to realize multiplexed ultrasound imaging to uniquely visualize locations of different species of gas vesicles. Mid-band fit spectral imaging technique demonstrated that stationary gas vesicles were efficiently localized in gel phantom and murine liver specimens by visualizing three-dimensional vessel structures. Clustered and unclustered gas vesicles were phagocytosed by murine macrophages to serve as carriers and beacons for the proposed multiplexed and single cell level imaging technique. The spatial distribution of macrophages containing clustered and unclustered gas vesicles were reconstructed by mid-band fit spectral imaging with pseudo-coloring scheme.

---

Fluorescent proteins such as green fluorescent protein (GFP) and red fluorescent protein (RFP) revolutionized biology and biochemistry^1^. Fluorescent proteins established a marker of gene expression and protein targeting in live cells and organisms. Successful development of ultrasound gene reporters for deep penetration and high resolution imaging will revolutionize deep tissue cell tracking with multiplexed imaging capability^2,3^. Similar to optical microscopies using GFP and RFP, ultrasound contrast agents that emit distinct spectrum under an ultrasound excitation may visualize the spatial distribution of different cell populations (Fig 1c, d). Hence, we discovered a new type of gas vesicles (GV) as ultrasound contrast agents and developed spectral imaging approach with pseudo-coloring to visualize the location of different types of GVs to test multiplexed imaging potential. We investigated multiplexed imaging potential using clustered GVs and high frequency ultrasound at millimeter scale using mid-band fit (MBF) spectral imaging. In principle, the implementation of multiplexed ultrasound MBF spectral imaging could bypass the need of and hence the issues with ex vivo fixation based imaging techniques and in vivo optical methods with optical windows to track cells that travel close to the surface. Proposed GV and MBF based imaging approach continuously images target cells and region for a long time because of the stability of GVs without the needs to break them.

**Figure 1.**
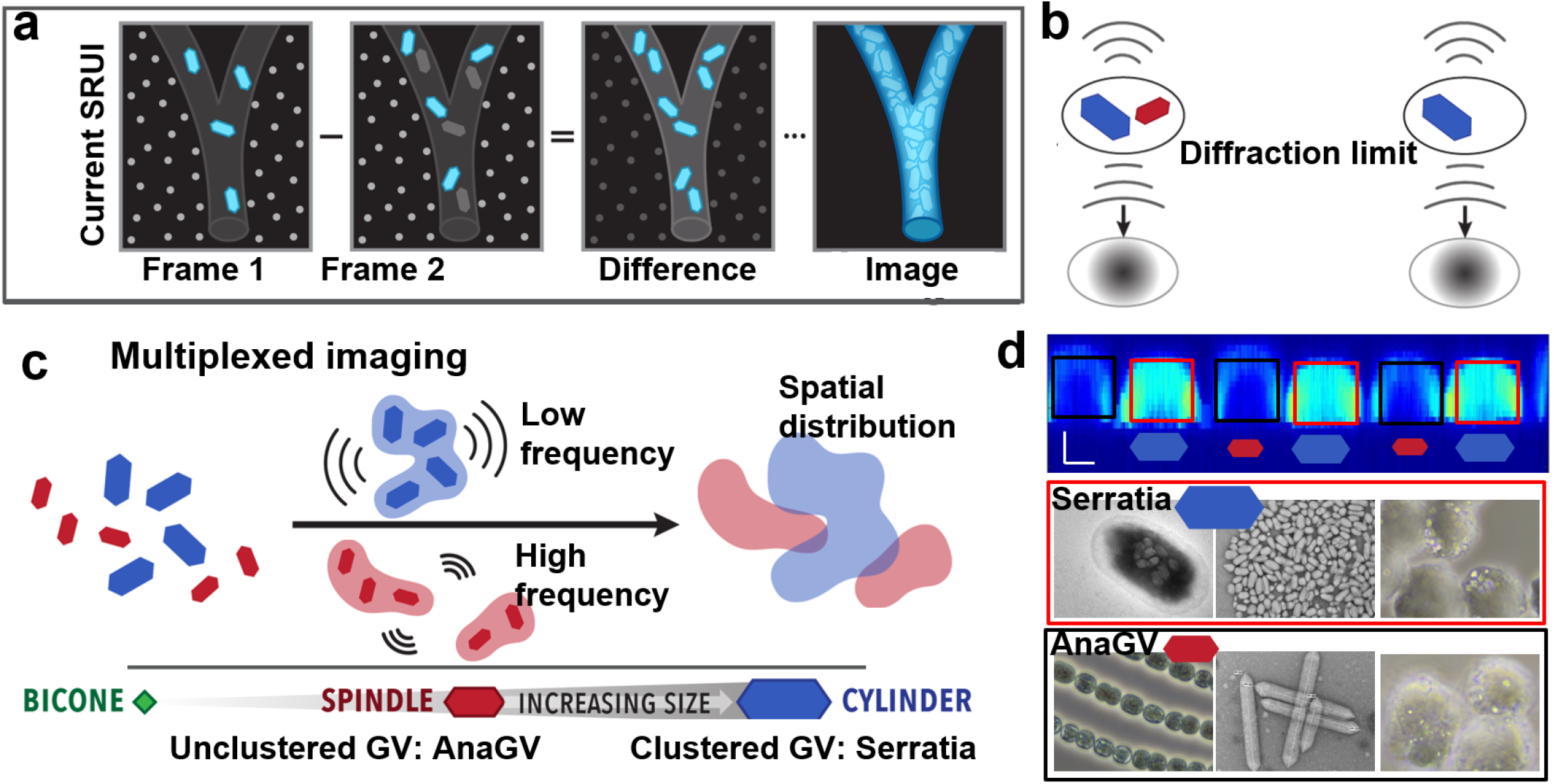
Multiplexed ultrasound spectral imaging of stationary gas vesicles (GV). **a.** Current superresolution ultrasound imaging (SRUI) subtracts consecutive frames to dampen background signal from surrounding tissues while increasing signals from moving microbubbles to visualize vessels. **b.** The spatial resolution is limited by the ultrasound system’s diffraction limit and the imaging acquisition time is limited by the concentration of microbubbles. Acquisition time and signal interference by diffraction limit are controversial and stationary and slowly moving object cannot be imaged by SRUI. **c.** Clustered GVs (blue, SerratiaGV) and unclustered GVs (red, AnaGV) emit distinct spectrum under ultrasound excitation by providing basis for multiplexed spectral imaging of stationary GVs. **d.** Clustered Serratia GVs and unclustered AnaGVs present distinct mid-band fit (MBF) signals. Serratia GVs are produced in rod-shape *Enterobacteriaceae* and AnaGVs are produced in *Anabaena flos-aquae.* Brighter and larger spots indicate clustered SerratiaGVs when they are internalized in RAW cells (red boxes). Scale bars: 1mm.

Unbreakable ground truths govern imaging modalities: 1) imaging depth and resolution are inversely proportional; and 2) resolution of an imaging system is bounded to about half the wavelength of sound waves or light, defined as the diffraction-limit. Revolutionizing a traditional perception of low spatial resolution and deep penetration intrinsic to ultrasound modality, modern ultrasound contrast agents (e.g. microbubbles [MB]), enable an unprecedented super-resolution ultrasound imaging (SRUI)^4,5^. The resolution of SRUI is directly related to the size of MB (less than 10 μm). Compared to the resolution of 8 MHz ultrasound waves with the penetration depth of 100 mm^6^, SRUI provides great improvement; 190 μm vs. 10 μm. Image reconstruction of SRUI uses subtraction between two consecutive frames of densely populated fast moving MBs (Fig. 1a). Subtraction between two frames identifies the MB location while the stationary tissue and background signals cancel out. The frame subtraction approach in SRUI is adapted from stimulated emission depleted (STED) microscopy by mimicking switching on fluorescence within confined area by overlapping stimulation light and donut-shaped deexcitation (or STED) light^7–9^. However, because the ultrasound signal from MB cannot be turned off, moving MBs are used to mimic signal flickering between two frames. A moving MB represents signals at two points of two frames. The main applications of SRUI have been the visualization of microvessels in the brain^5^ or kidney^10^ because of blood flow in vessels. The resolution of SRUI is related to the signal-to-noise ratio, bandwidth of backscattered echoes and the number of array elements used in the beamforming and does not follow ground truth #2 above. The Cramer-Rao lower bound determines the resolution limit of SRUI^11^. This SRUI has been a partial success to achieve SRUI with deep tissue capability. Stationary and slowly moving objects such as migrating cells and infiltrating cells within wound sites or tumor microenvironment cannot be imaged with current SRUI. Another requirement of SRUI is that MBs should be precisely located within focus using low concentration MB perfusion to avoid MB signal interference, which induces minutes-long acquisitions to fill the small vessels. For example, within 190 μm focus of 8 MHz ultrasound, only one MB is required for every subtraction between frames. Two or more MBs result in one blurred image due to the diffraction-limit of the ultrasound system (Fig. 1b). Densely populated MBs do not provide a true SRUI due to signal interference while sparse MBs increase imaging acquisition time significantly. Our group developed L1-homotopy based compressed sensing algorithm to improve the efficiency of the localization of point targets and imaging acquisition time^12^. However, usually, this strategy works fine for vessel visualization because edges and borders need to be identified instead of localizing each single MBs. The requirement of next generation SRUI is high resolution imaging capability of slowly moving and stationary targets.

We used MBF based spectral imaging approach for imaging stationary targets. MBF predicts the possibility of the existence of particles such as GVs and MBs by measuring spectral amplitude in dB of reflected waves from particles. Scattering from regular tissues and ultrasound contrast agents, GVs and MBs, present different MBF signal. MBF of backscattering from different sizes of GVs and background tissues were calculated and visualized by assigning pseudo-colors depending on MBF values to reconstruct a color map to visualize the locations of different types of GVs. To demonstrate the feasibility of GV and MBF based spectral imaging, we chose 130 MHz and 50 MHz ultrasound after considering the size and scattering pattern, which is usually determined by *ka* value, where *k* is wavenumber and *a* is the radius of a target under consideration. Small *ka* value indicates either small particle size or low ultrasound frequency^13^. Considering the sizes of GVs and the frequencies in consideration, *ka* ranges from 0.02 to 0.2 for 50 MHz and from 0.05 to 0.5 for 130 MHz ultrasound, respectively. In this diffusive scattering regime, the main source for ultrasound imaging is scattering by an ensemble of unresolved particles. In this regime, we speculated that diffusive scattering from individual and unclustered GVs and diffractive scattering from clustered GVs or GVs phagocytosed into cells may be differentiated to provide multiplexed images. In diffractive scattering, *ka* is close to 1, which is true for both 130 MHz and 50 MHz ultrasound reflected by clustered GVs, directivity is determined by the object shape, angle, and *ka*. The difference between clustered and unclustered GVs may provide clear multiplexed imaging source. Microbubbles (MBs) are developed for ultrasound contrast agents that contains gas core surrounded by lipid- or protein shells. MBs are FDA approved for disease diagnostic imaging sessions^14^. Due to their large size compared to cells and junction between connective tissues in vessels, MBs are restricted to the imaging of blood stream and increasing permeability of cells from outside^15^. MBs are physically unstable so they are diffused out within tens of minutes. Long-term imaging and extravasation and / or cell tracking have been limited to MB-enhanced ultrasound imaging technique. To address these challenges, nanometer-sized gas-filled protein structures that are produced by a range of bacteria and archaea are used. GVs are intracellular gas-filled protein structures that function as flotation devices to maintain a suitable depth in water^2–4,16–19^. GVs are perfect contrast agents for ultrasound due to high impedance mismatch. GVs are found in *cyanobacteria, halophilic archaea,* and *Bacillus megaterium* (Mega). *Baillus megaterium* gas vesicle protein (Gvp) was identified and successfully transferred to *E.co*li^20^. Mammalian cell expression was also demonstrated^21^. The wall of GVs is freely permeable to gas molecules and is composed of a small hydrophobic protein, GvpA, which forms a single-layer wall^22–25^. In addition, several minor structural accessory or regulatory proteins are required for GV formation^26^.

Serratia species are gram-negative and members of the Enterobacteriaceae. *Serratia* ATCC 39006 (Serratia) was first discovered at a salt marsh in New Jersey during a search for new antibiotic, antimicrobial, anti-cancer agents^27^. Serratia is the only member to produce GVs in the Enterobacteriaceae as a flotation organelle^28^. Genes required for GV production were sequenced as a 16.6 kb gene cluster encoding three distinct homologs of the main structural protein, GvpA^18,28–31^. While illuminating exciting area of multiplexed ultrasound imaging for deep tissue and higher resolution cell tracking using Serratia, we hope that the high genetic tractability of Serratia^18,28^ will enable faster research progress in GV biology for deep tissue multiplexed ultrasound imaging. Here, we uniquely combine clustered and unclustered GVs as ultrasound contrast agents and MBF spectral imaging for multiplexed imaging approach. We isolated a panel of gas vesicles from their host bacteria for multiplexed imaging that can emit distinct MBF spectrum. We demonstrated MBF spectral imaging to specifically visualize the location of clustered and unclustered GVs using 130 MHz and 50 MHz ultrasound when GVs are phagocytosed by murine macrophage. Foremost, by harnessing our high frequency ultrasonic transducers and ultrasound biomicroscopy (UBM) with GVs, we develop a general approach to realize multiplexed ultrasound imaging by visualizing the location of clustered and unclustered GVs. Moreover, by leveraging GV’s biocompatibility and stability, we internalized these GVs in phagocytic cells to serve as ultrasound imaging contrast agents that can be directly tracked with high frequency ultrasound at depth. Generated GVs were tested to visualize three-dimensional vessel structure of murine liver specimens ex vivo. Furthermore, multiplexed imaging capability was demonstrated that the locations of cells with clustered GVs and unclustered GVs were identified by distinct filter settings defined by MBF spectral signals similar to pseudo-coloring methods in fluorescence microscopy.

## Results

### Clustered Serratia GVs

Multiplexed ultrasound imaging may be achieved by imaging individualized and unclustered GVs and clustered GVs because their size differences can produce distinct echo spectrum. We isolated different types of bacteria including *Anabaena flos-aquae* (Ana), Mega, *Calothrix* sp. PCC 7601, and Serratia to find GVs with different morphology for multiplexed imaging. Gvp gene clusters of Serratia are well identified^18,28^. Isolated Serratia GVs (SGV) are naturally clustered (Fig. 2a, b) and have much lower aspect ratio compared to other GVs (Fig. 2b). While detailed Serratia culture conditions are described in Methods, we incubated Serratia in 20°C, 26°C, and 30°C culture media and measured Serratia proliferation and GV yield (GV expressing Serratia / all Serratia (%)) to find an optimized culture condition as shown in Fig. 2c. SGVs were synthesized in Serratia starting 13 hours of incubation at 26°C (C1 at Fig. 2c). GV expression was maximum when Serratia was incubated at 26°C (Fig. 2c). SGVs required at least 24 hours of incubation at 26°C to fully grow. Measured length and width were 237±36 nm and 128±12 nm, respectively (Supplementary Fig. 1a). Using this optimized culture conditions, we isolated SGV after 48 – 72 hours of incubation, which formed clustered and big flake shapes as big as 5 μm as shown in phase contrast images (PCI) and TEM images (CLUS, Fig. 2a, b). We tried urea, SDS, Triton X-100, Tween-20, Tween-80, and BSA to separate CLUS (UN) (details in Methods and Supplementary Fig. 1c). We chose urea due to it stability after treatment. Then, we tested cell cytotoxicity of CLUS and UN and GVs from Ana (AnaGV) and Mega (MegaGV). The cytotoxicity of GVs towards RAW 264.7 cells were assessed using the XTT cell proliferation assay. Only 20% of RAW cells survived after 2 hours of incubation with CLUS and UN (green and blue bars in Supplementary Fig. 1b). Prodigiosin and carbapenem are secondary metabolites produced by Serratia^29,31^. Prodigiosin is the red pigment (3 days CLUS in Supplementary Fig. 1b) and carbapenem induces apoptosis of cells, which leads significant cells death. Regular washing process did not remove prodigiosin and carbapenem and repeated washing process removes SGVs. Because the production of these secondary metabolites was related to quorum sensing and the production of GVs in Serratia^28^, we reasoned that long-term incubation may remove prodigiosin and carbapenem after the stable supply of oxygen and the Serratia colonies’ settlement near water-air interface. After one week of incubation, red Serratia culture media changed its color and after 2 weeks, culture media became yellow (Supplementary Fig. 1b). Isolated CLUS showed white cloud, which indicated compound changes after the long incubation (Supplementary Fig. 1b). XTT cell proliferation assay indicated that cell cytotoxicity to RAW cells of CLUS after two to three weeks of incubation is similar level to control group (CTL) as shown in Supplementary Fig. 1b. T his confirms that long-term incubation excludes these secondary metabolites from Serratia. We also observed that SGV and AnaGV were intact after an overnight incubation at 37°C with 5% CO_2_ (Supplementary Fig. 1d). We tested multiplexed potential of CLUS and other GVs such as UN, AnaGV, and MegaGV in the following section.

**Figure 2.**
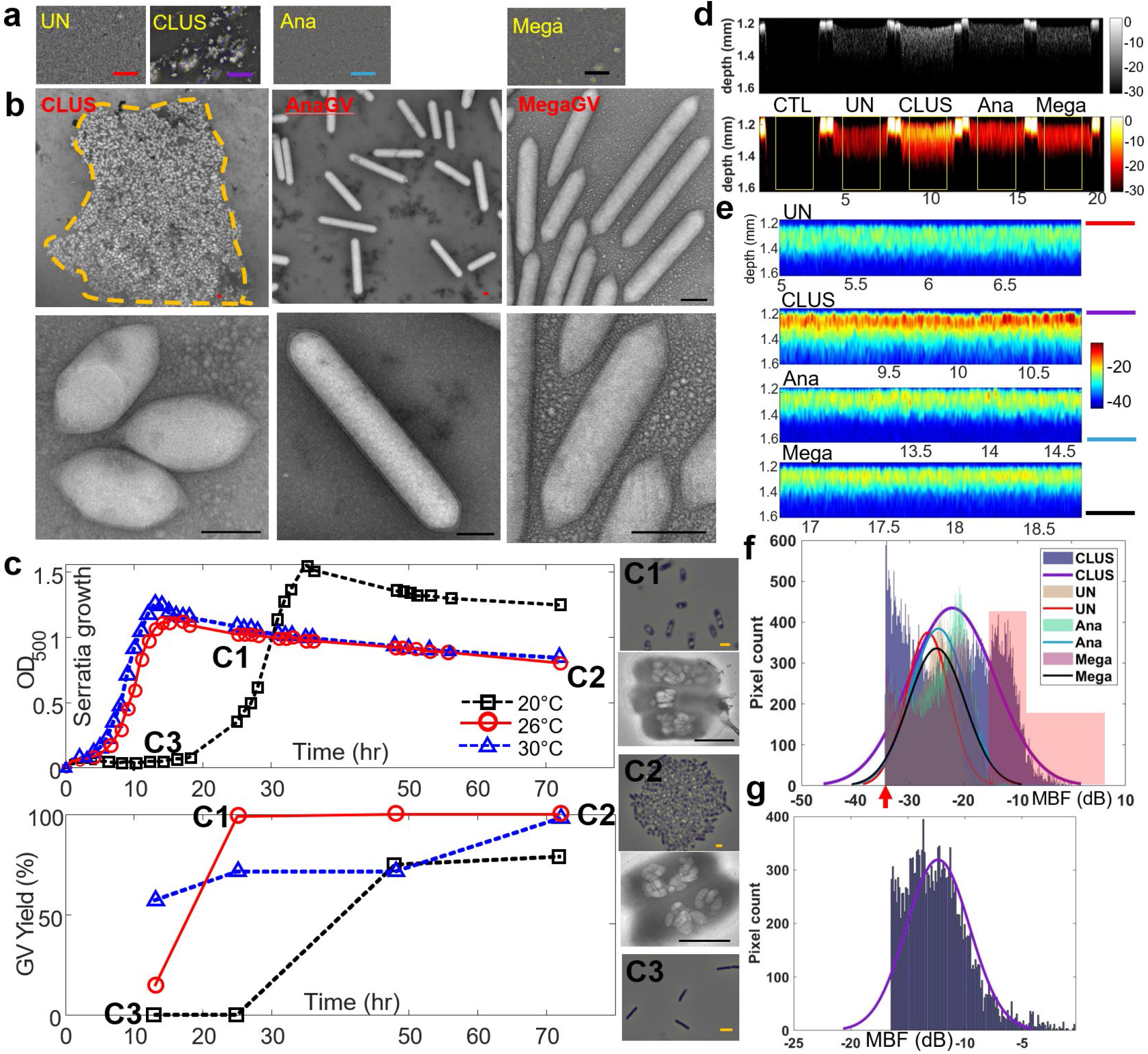
Clustered GVs from *Serratia.* **a.** Phase contrast images (PCI) of unclustered GVs (UN) and clustered GVs (CLUS) from *Serratia* and GVs from *Anabaena flos-aquae* (Ana) and *Bacillus megaterium* (Mega) show large GV clusters of CLUS. Salt and pepper patterns indicate unclustered GVs including UN, Ana, Mega in micrometer scale. Scale bars: 10 μm. **b.** TEM images of CLUS, AnaGVs, and MegaGVs show detailed structure of GVs. CLUS form a large cluster (dashed line) that cannot be dissolved by external flows or centrifugation. Scale bars: 100 μm. **c.** To isolate GVs from *Serratia*, we optimized culture condition by changing temperature and incubation time. Optimized temperature is 26°C (red circles) and maximum GV production in Serratia required at least 24 hours of incubation (C1) to reach 100% GV yield. PCI images and TEM images shows *Serratia* and GVs in their cytoplasm at (C1) 24 hrs of incubation at 26°C, (C2) 72 hrs of incubation at 26°C, and (C3) 13 hrs of incubation at 30°C, Black cylindrical shapes in PCI indicate *Serratia* and bright dots indicate GVs in *Serratia.* TEM images shows individual GVs produced in *Serratia.* Scale bars: 500 nm. **d.** B-mode and MBF spectral images of control (CTL), UN, CLUS, Ana, Mega GVs scanned by 130 MHz ultrasound. GVs are mixed with 1% gelatin at the final OD_500_ of 10. **e.** Zoom-in MBF spectral images of yellow boxes in (**d**) indicates that CLUS signal is distinct from MBF signals from other GVs. **f.** Histogram of MBF signal of all pixels in yellow boxes shows that CLUS signal has distinct MBF signal indicated by red regions.

### Multiplexed ultrasound imaging using CLUS

We developed 130 MHz and 50 MHz ultrasonic transducers and measured performance to investigate multiplexing potential of different morphology of GVs (details in Methods and Supplementary Fig. 2). A 1% gel was mixed with UN, CLUS, AnaGV, MegaGV for the final GV concentration at OD_500_ of 10 (Supplementary Fig. 2c). The phantom was scanned by ultrasound biomicroscope (UBM) with 1μm step size for 20 mm along width direction (Supplementary Fig. 2c). A B-mode on top row and MBF spectral image on bottom row are shown in Fig. 2d. Yellow regions are zoomed to visualize MBF signal distribution of UN, CLUS, AnaGV, and MegaGV (Fig. 2e). All MBF signals were normalized by the maximum MBF signal within four GV regions. CLUS MBF signals are stronger than other GV MBF signals (Fig. 2e, f). MBF signals smaller than mean of background (CTL) + 2 * standard deviation of CTL was truncated (red arrow in Fig. 2f) because this is basal level signal. To compare MBF signals of UN, CLUS, AnaGV, and MegaGV, the average and standard deviation were measured using all pixels within yellow boxes in Fig. 2d and measured to be −26.6 ± 4.0 dB (UN), −22.1 ± 7.9 dB (CLUS), −24.7 ± 5.1 dB (AnaGV), and −25.0 ± 5.2 dB (MegaGV). The main difference in histogram in Fig. 2f are red rectangular regions, where most pixel counts are from CLUS. To differentiate MBF signal from clustered GV and unclustered GV, we took mean of MBF of UN / AnaGV / MegaGV + 2* standard deviation of MBF of UN / AnaGV / MegaGV. The mean of these values was −16.5 dB, which was used to truncate CLUS MBF signal shown in Fig. 2g. Mean and standard deviation of this truncated CLUS MBF signal in Fig. 2g is −12.4 ± 2.7 dB. From the experiment, CLUS MBF signal (CMS) and individualized GV’s MBF signal (IMS) including UN, Ana, and Mega can be defined to test multiplexing potential of GV based MBF spectral imaging. One possible combination of CMS and IMS are −17.8 dB (−12.4 −2*2.7) and −34.6 dB (−26.6 – 2*4.0), which can serve as filters to identify locations of CLUS and UN. From this multiplexing strategy, MBF signals above CMS can be considered as clustered GV signal and MBF signals between CMS and IMS can be considered as unclustered GV signal. We also tested this multiplexed imaging potential using 50 MHz ultrasound (Supplementary Fig. 3). However, in this experiment, because OD_500_ is too high and GVs are densely packed, resulting in meaningless MBF signal distribution, we decided to reduce concentration of GVs and apply multiplexed imaging approach by defining CMS and IMS.

### Multiplexed ultrasound imaging using sparsely distributed GVs

For multiplexed MBF spectral imaging, the thresholds of CMS and IMS were investigated using decreased concentration of GVs at OD_500_ of 0.022, 0.044, and 0.088. The hypothesis is that clustered GVs emit distinct MBF spectrum, which can be used to differentiate clustered GVs and individual and unlcustered GVs. We scanned five wells that contains CTL, UN, CLUS, Ana, Mega and post processed radiofrequency (RF) data to acquire MBF signals (Supplementary Fig 2c). Two independent image scans were performed, and two samples were scanned per one independent image scan at three OD_500_. Figure 3 represents MBF spectral imaging from one experiment. MBF spectral images and their zoom-in images at OD_500_ of 0.022 (Fig. 3a, c) and 0.088 (Fig. 3e, g) show the distribution of GVs depending on GV concentration of UN, CLUS, Ana, and Mega. Mean and standard deviation of MBF signals within yellow boxes are shown in Figures 3b, f. The number of detected GV locations at these concentrations are approximately 20 or less. Each detected GV included approximately 40 pixels. Within yellow boxes of each well containing no GVs (CTL), UN, CLUS, Ana, and Mega, 800 pixel values were used to calculate mean and standard deviation of MBF signals. MBF of CLUS for OD_500_ of 0.022 and 0.088 are −18.5 ± 4.5 dB and −21.1 ± 1.3 dB, respectively. CLUS MBF is clearly distinct from other GV’s MBF, which are ranging from −31 to −27 for OD_500_ of 0.022 and from −27 to −24 for OD_500_ of 0.088 (Fig. 3b, f). CMS was assigned by calculating CLUS MBF – 2*standard deviation of CLUS MBF. For OD_500_ of 0.022, CMS was −27.5 dB and for OD_500_ of 0.088, CMS was −23.7 dB. IMS was defined as the mean of UN / Ana / Mega MBF – 2*standard deviation of UN / Ana / Mega MBF. For OD_500_ of 0.022, IMS was −36.4 dB and for OD_500_ of 0.088, IMS was – 30.6 dB. OD_500_ of 0.044 MBF spectral imaging was also performed. To define CMS and IMS to cover OD_500_ between 0.022 and 0.088, mean values of CMS and IMS of three OD_500_ were calculated using four different measurements (Fig. 3j), where clear distinction between CLUS MBF signal in a red rectangle and other GV MBF signals in a yellow rectangle.

**Figure 3.**
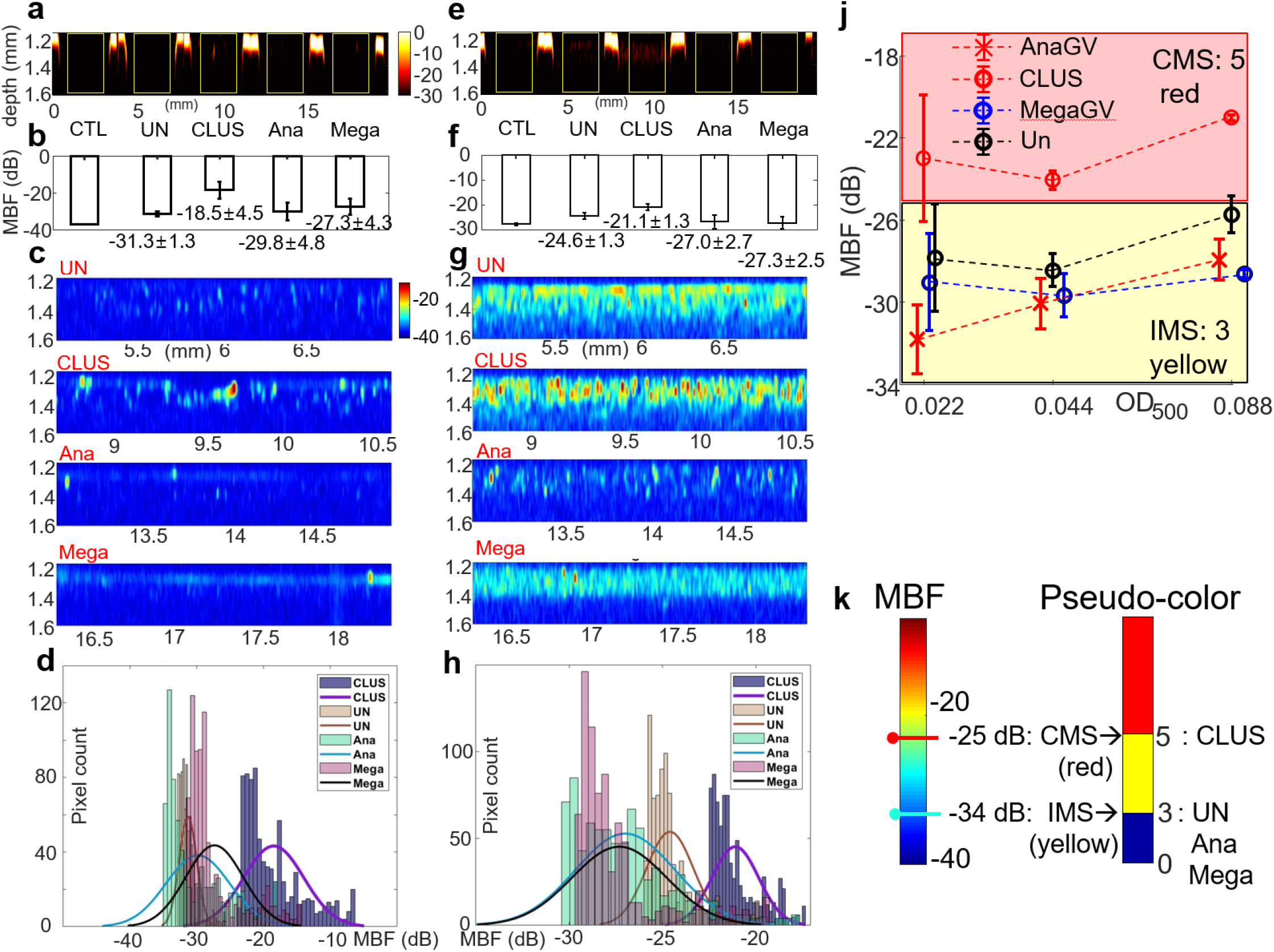
MBF spectral images of sparsely distributed GVs. MBF spectral images of CTL, UN, CLUS, Ana, and Mega at OD_500_ of (**a**) 0.022 and (**e**) 0.088 and zoom-in images of UN, CLUS, Ana, and Mega at OD_500_ of (**c**) 0.022 and (**g**) 0.088 show the location of detected GVs by 130 MHz ultrasound. Using 800 highest MBF pixels in (**c**) and (**g**), the mean and standard deviation of MBF signals from UN, CLUS, Ana, and Mega were calculated at OD_500_ of (**b**) 0.022 and (**f**) 0.088. **d.** Histogram of MBF signals from UN, CLUS, Ana, and Mega at OD_500_ of (**d**) 0.022 and (**h**) 0.088 presents clear distinction between CLUS and other unclustered GVs. **j.** The mean and standard deviation of MBF signals from Ana, CLUS, Mega, UN were measured at OD_500_ of 0.022, 0.044, and 0.088 (n=4). CMS was defined by calculating the mean of CLUS MBF – 2*standard deviation of CLUS MBF at three OD_500_. The mean of CMS at OD_500_ of 0.022, 0.044, and 0.088 was first calculated and their mean was defined as CMS to detect CLUS signals at OD_500_ of 0.022-0.088. Similarly, IMS was defined by taking the mean of IMS of UN, Ana, and Mega. **k.** MBF images were converted into two-color multiplexed images to differentiate the location of CLUS and unclustered GVs including UN, Ana, and Mega. CMS and IMS values defined was used to assign pseudo-colors. Pixels with MBF signals over CMS were assigned red color and pixels with MBF signals between CMS and IMS were assigned yellow. Red pixels indicate the locations of CLUS and yellow pixels indicate the locations of unlcustered GVs (UN, Ana, and Mega).

Pseudo-coloring was achieved by assigning value 5 with red pseudo-color if MBF signal is above CMS and value 3 with yellow pseudo-color if MBF signal is between CMS and IMS (Fig. 3j, k). This is similar to pseudocolor scheme in fluorescence microscope. We expected to differentiate clustered GV locations and unclustered GV locations. We mixed CLUS and UN (C+U), CLUS and Ana (C+A), and CLUS and Mega (C+M) to investigate potential multiplexed imaging capability using MBF spectral imaging with CMS and IMS. OD_500_ of each GV species was 0.022 and 0.044, resulting in total concentrations of OD_500_ of 0.044 and 0.088. UBM system were used to scan GV mixture phantom (Supplementary Fig. 2b, c). MBF spectral and B-mode images of total OD_500_ of 0.044 and 0.088 cases were reconstructed (Fig. 4a, d and 4e, g). Multiplexed images after applying CMS and IMS based pseudo-coloring were obtained (Fig. 4c, h). Red regions predict the location of CLUS and yellow regions predict the locations of UN / Ana / Mega.

50 MHz ultrasound could not detect individual and clustered GVs at OD_500_ of 0.022 – 0.088. Such a low concentration, GVs are too small compared to the wavelength of 50 MHz. Because realizing a deep tissue and single cell level resolution is the goal of proposed approach, we changed imaging targets from individual and clustered GVs to single cells with GVs as ultrasound contrast agents. The size of single cells is comparable to the wavelength of 130 MHz and 50 MHz ultrasound. We hypothesized that if GVs are internalized in the cytoplasm of cells, then echo signals and MBF spectral signals may increase and be easily distinguished between UN, CLUS, Ana, and Mega GVs. We tested this using RAW 293 cells (ATCC).

**Figure 4.**
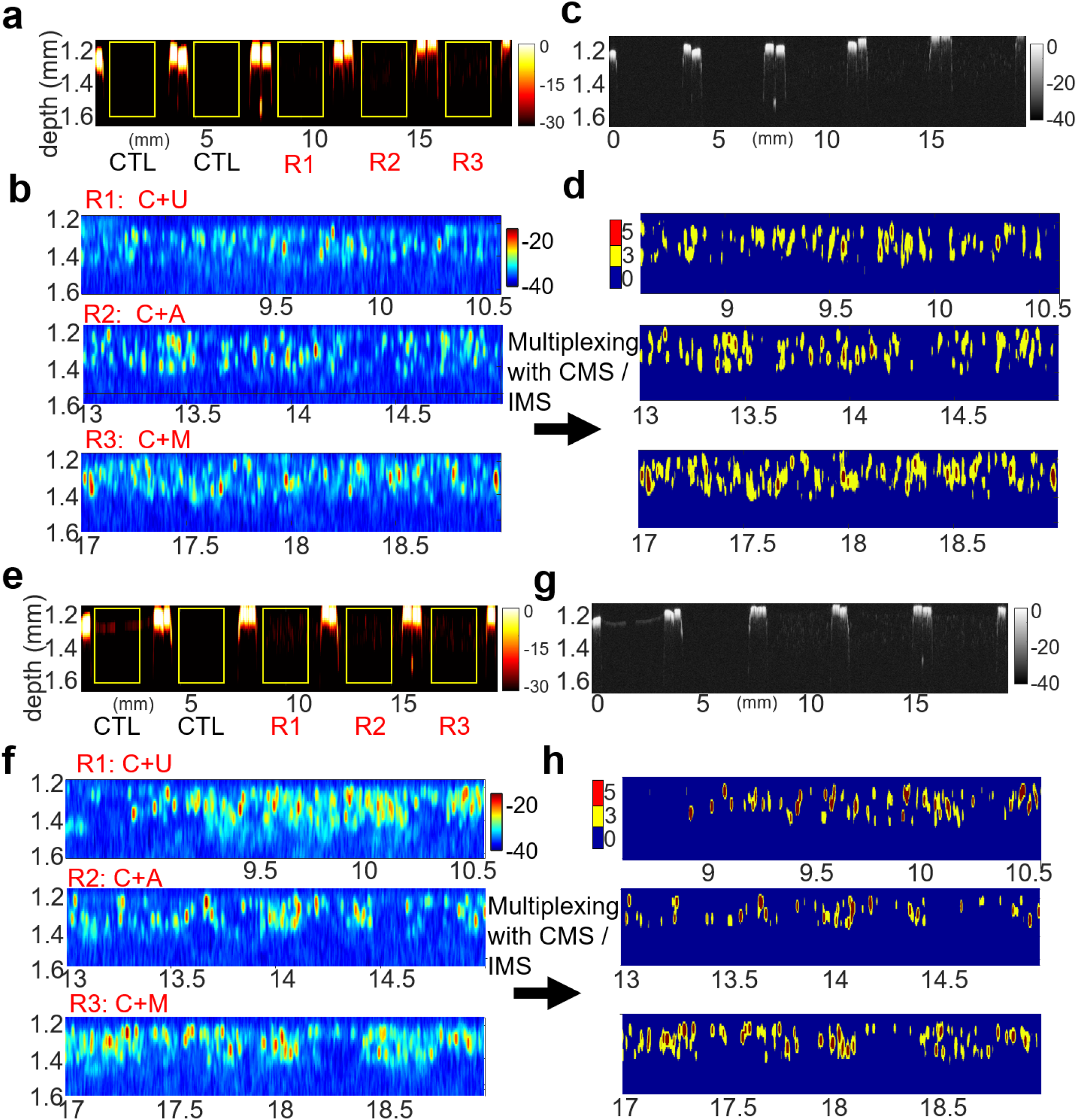
Multiplexed MBF spectral images of mixed GVs. **(a, c, e, g)** First two regions are 1% gel phantom without GVs as CTL**. (a, e)** MBF spectral images and **(c, g)** B-mode images show overall distribution of mixed GVs. **(a, c)** Regions R1, R2, and R3 were 1% gel phantoms mixed with CLUS at OD_500_ of 0.022 and UN at OD_500_ of 0.022 (C+U), CLUS at OD_500_ of 0.022 and Ana at OD_500_ of 0.022 (C+A), and CLUS at OD_500_ of 0.022 and Mega at OD_500_ of 0.022 (C+M). **b.** Zoom-in MBF spectral images of R1, R2, and R3 and **d.** multiplexed images using pseudo-color approach with CMS and IMS represent the locations of CLUS and unclustered GVs. **(e, g)** Regions R1, R2, and R3 were 1% gel phantoms mixed with CLUS at OD_500_ of 0.044 and UN at OD_500_ of 0.044 (C+U), CLUS at OD_500_ of 0.044 and Ana at OD_500_ of 0.044 (C+A), and CLUS at OD_500_ of 0.044 and Mega at OD_500_ of 0.044 (C+M). **f.** Zoom-in MBF spectral images of R1, R2, and R3 and **h.** multiplexed images using pseudo-color approach using CMS and IMS represent the locations of CLUS and unclustered GVs.

### Visualization of vessels in murine liver using AnaGV and CLUS

Before we used RAW cells to test multiplexing, we imaged vessels in mouse liver specimens. Mouse liver was perfused and injected with 1 ml of AnaGVs and SGVs at OD_500_ of 20. B-mode, MBF, and filtered MBF (fMBF) images were reconstructed (Fig. 5a-c, and 5e-g). MBF images with psuedo-color is defined as fMBF. Here, MBF values over CMS was assigned white and MBF values between CMS and IMS were assigned orange color. Total scan areas of AnaGV and CLUS injected liver specimens were 5 mm along the width direction by 360 μm along elevation (z) direction and 5 mm by 260 μm, respectively (Figs. 5d, h and Supplementary Fig. 2c, 4). Scanning step sizes along the width direction and z-direction were 1 μm and 5 μm, respectively. Later by stacking 2D images along z-direction, 3D rendering was created to visualize vessels (Figs. 5d, h, Supplementary Videos 1, 2). B-mode and MBF spectral images show cross-sectional views of vessels visualized by AnaGVs (Red arrow in Figs. 5a, b) and CLUS (Red arrow in Fig. 5e, f). These images were 23^th^ section, 110 μm from the origin along z-direction for AnaGV injected liver and 38^th^ section, 185 μm from the origin along z-direction for CLUS injection liver (Figs. 5d, h). Zoom-in images of B-mode and MBF spectral images of red box regions in Figs. 5a, b and 5e, f represent the location of vessel in the cross-section. Vessel signals due to GVs and tissue scattering were filtered by CMS and IMS. CMS was defined as the mean of MBF signal of vessel regions – 2*standard deviation of MBF signal of vessel regions. IMS were defined as the mean of MBF signal of tissue regions – 2*standard deviation of MBF signal of tissue regions. MBF spectral images in Figures 5b and 5f were filtered with CMS and IMS by assigning white pseudo-color if MBF signals were larger than CMS and orange pseudocolor if MBF signals were between CMS and IMS values. fMBF images efficiently rejected tissue signals from out of focus area and emphasized vessel location at focus as shown in Figures 5c and 5g. 3D rendering using 73 sections for AnaGVs injected liver and 53 sections for CLUS injected livers were generated using fMBF spectral images (Figs. 5d, h). Vessel structures are visualized as white curved pipe-shaped regions in 3D rendering in Figures 5d and 5h. Red arrows in Figs. 5a, c, d and 5e, g, h indicate the same vessel crosssections. Rotational video clips show reconstructed 3D rendering of these two mouse liver specimens (Supplementary Videos 1, 2). The hematoxylin-eosin (H&E) histology images of the same mouse liver specimens show cross-sectional views of vessels indicated as hollow holes (Supplementary Fig. 4). **Multiplexed imaging of RAW cells with GVs.** We incubated UN, CLUS, Ana, and Mega with RAW cells (UN_RAW_, CLUS_RAW_, Ana_RAW_, and Mega_RAW_) for two hours after flipping dishes to promote phagocytosis of RAW cells.

**Figure 5.**
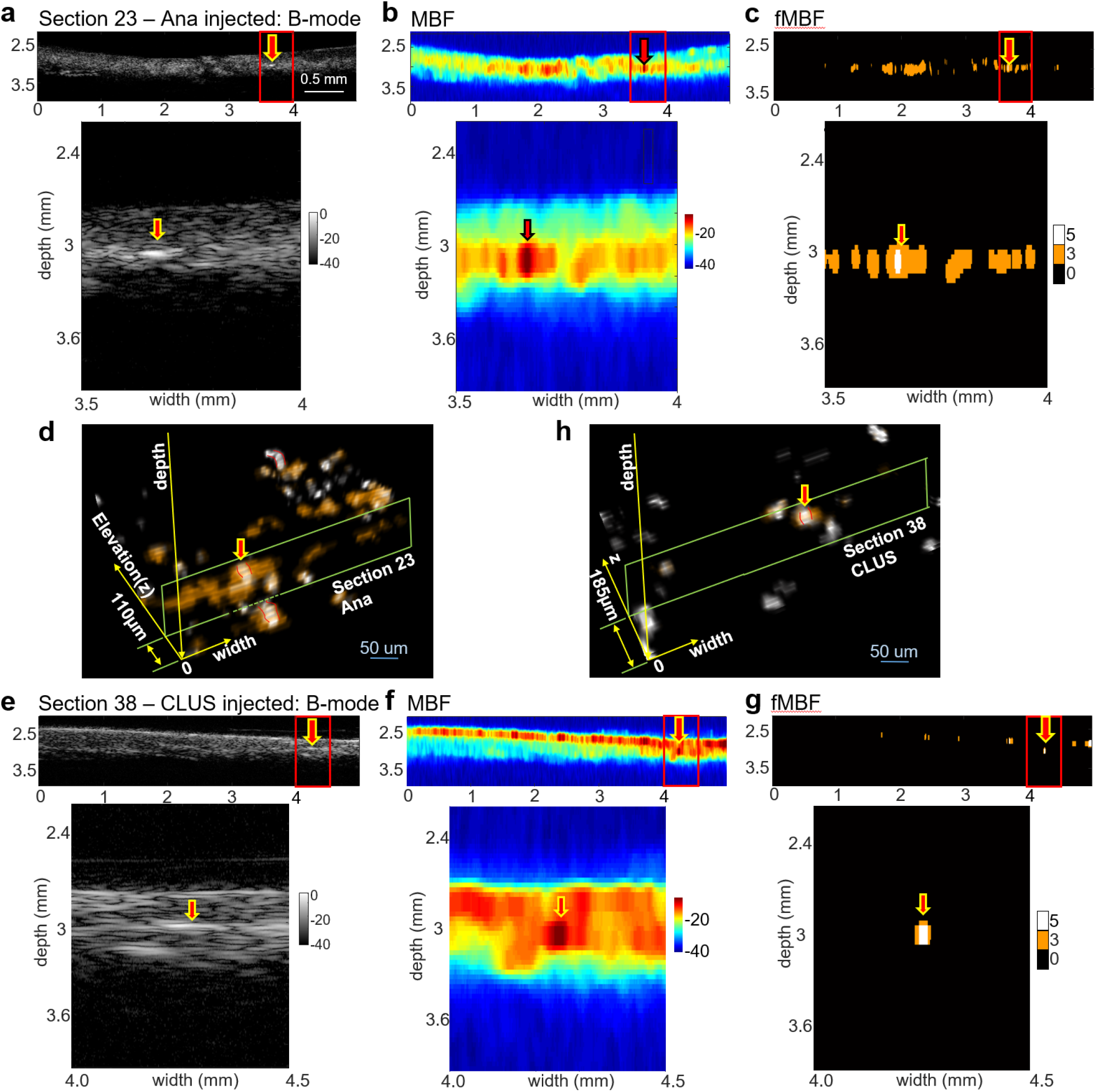
Mouse liver vessel visualization using AnaGV and CLUS. **a.** B-mode, **b.** MBF spectral image, and **c.** filtered MBF (fMBF) spectral images using CMS and IMS values visualize the location of a vessel injected with AnaGVs. **d.** Three-dimensional rendering by stacking fMBF images with 5 μm step size along elevation (z) direction shows three-dimensional vessels in liver specimen as indicated by white curved lines surrounded by red lines. Images in (**a, b, c**) are the 23th section which is 110 μm away from the front surface. **e.** B-mode, **f.** MBF spectral image, and **g.** filtered MBF (fMBF) spectral images using CMS and IMS values visualize the location of a vessel injected with CLUS. **h.** Three-dimensional rendering by stacking fMBF images with 5 μm step size along elevation (z) direction shows three-dimensional vessels in liver specimen as indicated by white curved lines surrounded by red lines. Images in **e. f. g.** are the 38th section which is 185 μm away from the front surface.

We measured MBF signal from 0.2 million (M), 0.5 M, 1 M, and 2 M cells without GVs. We chose 0.2 M and 0.5 M number of cells because these two cell numbers did not show significant MBF signal increase (Supplementary Fig. 5). To test signal detection limit, we also tested 50000 cells. We calculated a volume ratio of cells within a focus of 130 MHz and 50 MHz ultrasound. Within a voxel of 2 mm (width direction) X 400 μm (depth) X 10 μm (z-direction) for 130 MHz and 2 mm X 600 μ X 30 μm for 50 MHz, each RAW cell with GVs were expressed by approximately 40 pixels. Each RAW cell can be modeled as a sphere with the diameter of 10 μm. If 0.5 M cells are mixed with 60 μl PBS and 1% gel, then the volume ratios of one RAW cell in 60 μl gel phantom is 0.436% 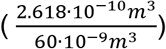. If 0.2 M and 50000 cells are mixed, the volume ratio to gel phantom is 0.175% and 0.044%, respectively. Total number of pixels were 382000 to reconstruct MBF spectral images of 2 mm X 400 μm region for 130 MHz ultrasound by multiplying 2000 (width direction) and 191 (depth) pixels. Considering the probability of one RAW cell detection (0.436%) within 2000 by 191 pixels results in approximately 42 cells in MBF spectral images. By multiplying 42 cells and 40 pixels per each RAW, 1620 pixels express RAW cells within 2000 by 191 MBF spectral images if 0.5 M cells are used. We calculated mean and standard deviations of MBF using 2000 pixels, 800 pixels, and 200 pixels for 0.5 M, 0.2 M, and 50000 RAW cells based on volume ratio of RAW cells within imaging slices. For 50 MHz ultrasound, we used 3000 pixels, 1200 pixels, and 300 pixels after considering an increased size of one imaging slice.

For an initial test, we measured MBF signal of 0.5M UN_RAW_, CLUS_RAW_, Ana_RAW_, and Mega_RAW_ using 2000 pixels based on volume ratio of cells (Fig. 6a-c). Obtained MBF signals were −21.4 ± 2.9 dB for UN_RAW_, −13 ± 3.0 dB for CLUS_RAW_, −29 ± 3.0 dB for Ana_RAW_, and −23.8 ± 3.0 dB for Mega_RAW_, respectively (Fig. 6a, b). Zoom-in images of yellow box regions in Fig. 6a show a detailed location of RAW cells with GVs (Fig. 6b). The histogram of MBF pixel values of UN_RAW_, CLUS_RAW_, Ana_RAW_, and Mega_RAW_ indicates distinct MBF signals from CLUS_RAW_ (Fig. 6c). We confirmed multiplexed imaging potential by comparing MBF signal and images of RAW cells taken by phase contrast imaging (PCI, Fig. 6e), confocal microscopy (Fig. 6f, g), and TEM (Fig. 6h). Large bright dots in PCI images and green fluorescent protein (GFP) signals in confocal microscopy images indicate phagocytosed CLUS in RAW cells (yellow arrows in Fig. 6e-g). TEM images in Fig. 6h confirm clustering of CLUS in RAW cells by comparing other GV internalization patterns in RAW cells (Supplementary Fig. 6).

**Figure 6.**
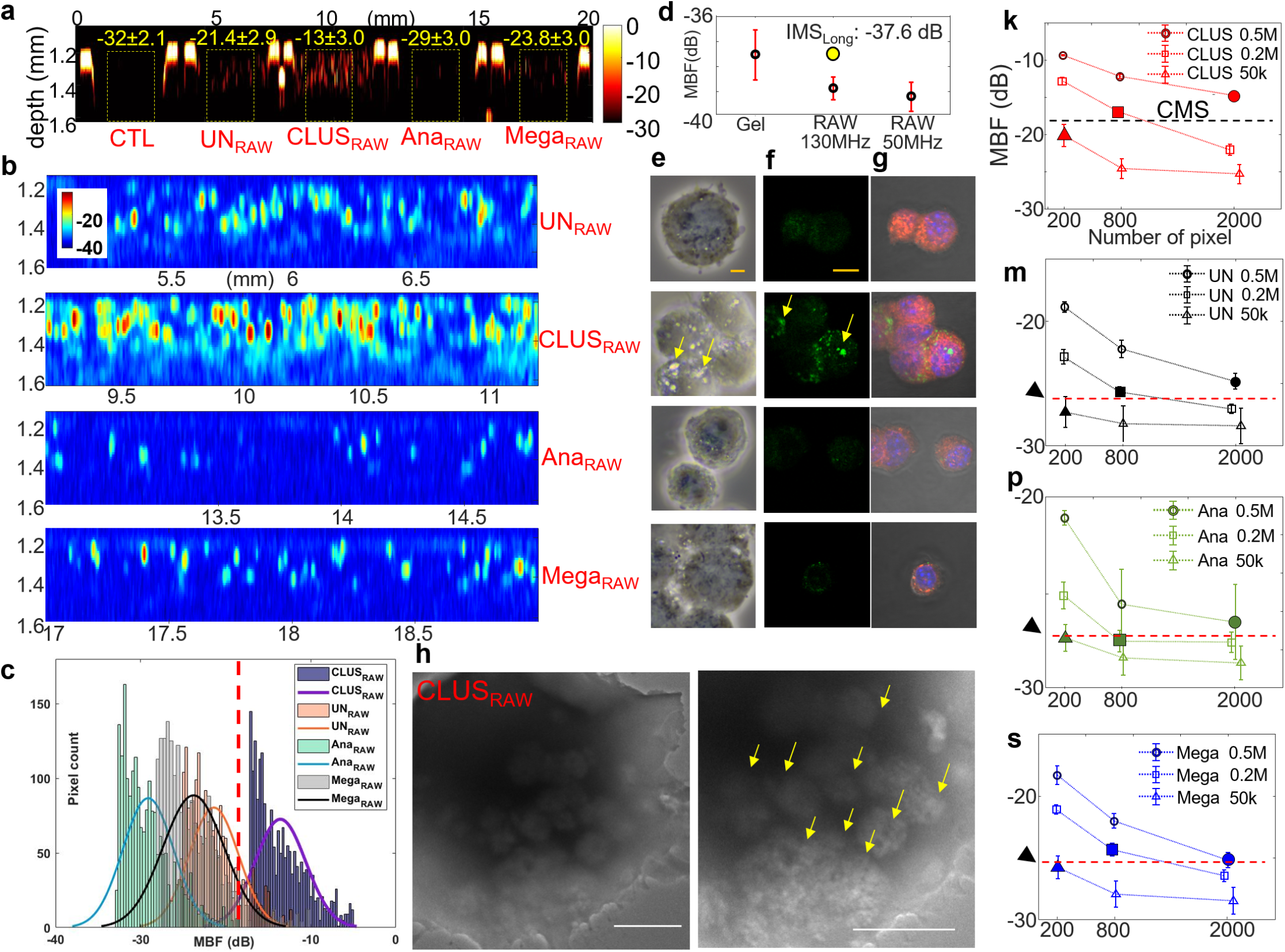
Single cell multiplexing. **a.** MBF spectral images of RAW cells (CTL) and RAW cells with UN (UN_RAW_), CLUS (CLUS_RAW_), Ana (Ana_RAW_), and Mega (Mega_RAW_). Means and standard deviations of MBF signal from CLUS_RAW_ are distinct from UN_RAW_, Ana_RAW_, Mega_RAW_., which shows distinct pixel counts in (**c**) the histogram. **b.** Zoom-in MBF spectral images of UN_RAW_, CLUS_RAW_, Ana_RAW_, and Mega_RAW_ indicate the distribution of GVs internalized by RAW cells. **d.** Basal levels (CTL) of MBF signal of 1% gel phantom, 0.5M, 0.2M, and 50000 RAW cells without GVs imaged by 130 MHz ultrasound and 50 MHz ultrasound were measured. IMS_Long_ was calculated by mean of CTL MBF + 3*standard deviation of MBF of CTL to reconstruct multiplexed images. **e.** PCI images and **f-g.** confocal images of UN_RAW_, CLUS_RAW_, Ana_RAW_, and Mega_RAW_ indicate the distribution of GVs in cytoplasm of RAW cells. Bright dots in PCI and GFP in confocal images indicate GVs. CLUS shows stronger and larger signals than other GVs, which confirms the clustered and larger GV size in RAW cells. **h.** TEM confirms clustering of CLUS in RAW cells (yellow arrows). TEM images of UN, Ana, and Mega in Supplementary Fig. 6. shows unclustered GV pattern in RAW cells. **k.** CMS for multiplexed imaging was defined by taking the mean of MBF signal from 0.5M, 0.2M, 50000 CLUS_RAW_ using 2000, 800, and 200 pixels (solid circle, square, and triangle) by considering the number of cells within the imaging slice. Similarly, IMS_short_ was defined by the mean of means of MBF signals (arrow head and red dashed lines) from **m.** UN_RAW_, **p.** Ana_RAW_, and **s.** Mega_RAW_. Error bars indicate ± one standard deviation.

MBF values for CMS and IMS were developed for multiplexed imaging. Based on histogram (Fig. 6c), CMS was defined by taking the mean of CLUS_RAW_ – 2* standard deviation of CLUS_RAW_ (Fig. 6k). MBF signals larger than this CMS was considered as clustered GV signals. IMS was defined as the mean of [UN_RAW_, Ana_RAW_, Mega_RAW_] – 2* standard deviation of [UN_RAW_, Ana_RAW_, Mega_RAW_] (IMS_short_) (Fig. 6m, 6p, 6s). Another IMS value was defined the mean of MBF of CTL + 3*standard deviation of MBF of CTL (IMS_Long_). IMS_short_ and IMS_Long_ corresponds short exposure and long exposure time in fluorescence microscope. IMS_short_ rejects larger MBF signals than IMS_Long_ does. To set up CMS, IMS_short_, and IMS_Long_, we performed two independent experiments with two samples per each experiment for 0.5M cells and three independent experiments with two samples per each experiment for 0.2M and 50000 cells. Basal level (MBF of CTL) was measured to be −38.8 ± 0.4 dB by calculating the mean and standard deviation of MBF signal using 0.5M, 0.2M, and 50000 RAW cells (RAW 130 MHz in Fig. 6d). IMS_Long_ was calculated to be −37.6 dB (yellow dot Fig. 6d). CMS was calculated to be −18.5 dB by taking the mean of MBF signals of 0.5M, 0.2M, 50000 CLUS_RAW_ using 2000, 800, and 200 pixels (solid circle, square, and triangle in Fig. 6k). The mean of MBF of 0.5M, 0.2M, 50000 UN_RAW_, Ana_RAW_, Mega_RAW_ (solid circles, squares, and triangles in Fig. 6m, 6p, 6s) was −26.9 dB, −28.7 dB, −25.5 dB as indicated black arrow heads and dashed red lines in Fig. 6m, 6p, 6s. IMS_short_ was calculated to be −27 dB by taking the mean of these values. These values were used to apply multiplexed imaging of RAW cells with clustered and unclustered GVs.

We performed the same experiments using 50 MHz ultrasound. CLUS_RAW_ exhibits distinct MBF signal compared to other GVs in RAW cells as shown in Supplementary Fig. 7. Lateral resolution of 50 MHz ultrasound is 30 μm, which shows blurred MBF images for CLUS_RAW_. (Supplementary Fig. 7b).

### Multiplexed MBF imaging by mixing RAW cells with GVs

We used pre-determined CMS as −18.5 dB, IMS_short_ as −27 dB, and IMS_Long_ was −37.6 dB. A mixture of Raw cells with CLUS and UN (CLUS+UN_RAW_), CLUS and Ana (CLUS+Ana_RAW_), and CLUS and Mega (CLUS+Mega_RAW_) were imaged to test multiplexed spectral imaging potential. The total number of cells after mixing were 0.5M and 0.2M cells. Figure 7 represents multiplexed MBF spectral imaging of 0.5M cells. The first image in Figures 7b–7d are zoom-in MBF spectral images of yellow dashed region in Fig. 7a. Pixel values over CMS were assigned red pseudo-color and pixels between CMS and IMS values were assigned yellow pseudo-color shown in the second and the third rows in Figures 7b–7d. Red pseudo-color indicates the existence of RAW cells with CLUS and yellow pseudo-color indicates the existence of RAW cells with unclustered GVs such as UN, Ana, or Mega. Images rendered with IMS_Long_ has much larger yellow regions than images with IMS_short_ because IMS_Long_ includes pixels with smaller MBF values. The histograms in Figs. 7b, 7c, 7d indicate that RAW cells with GVs emit higher MBF spectral signals compared to CTL. Red vertical lines, blue lines, and blue dashed lines in the histograms indicate CMS, IMS_short_ and IMS_Long_ values. Different pseudo-coloring using IMS_short_ and IMS_Long_ is analogous to changing exposure time in fluorescence microscope. Multiplexing of 0.2M cells with 130 MHz ultrasound and 0.2M and 0.5M cells with 50 MHz ultrasound demonstrate that clustered GVs and unclustered GVs can be differentiated by the proposed method (Supplementary Fig. 8).

**Figure 7.**
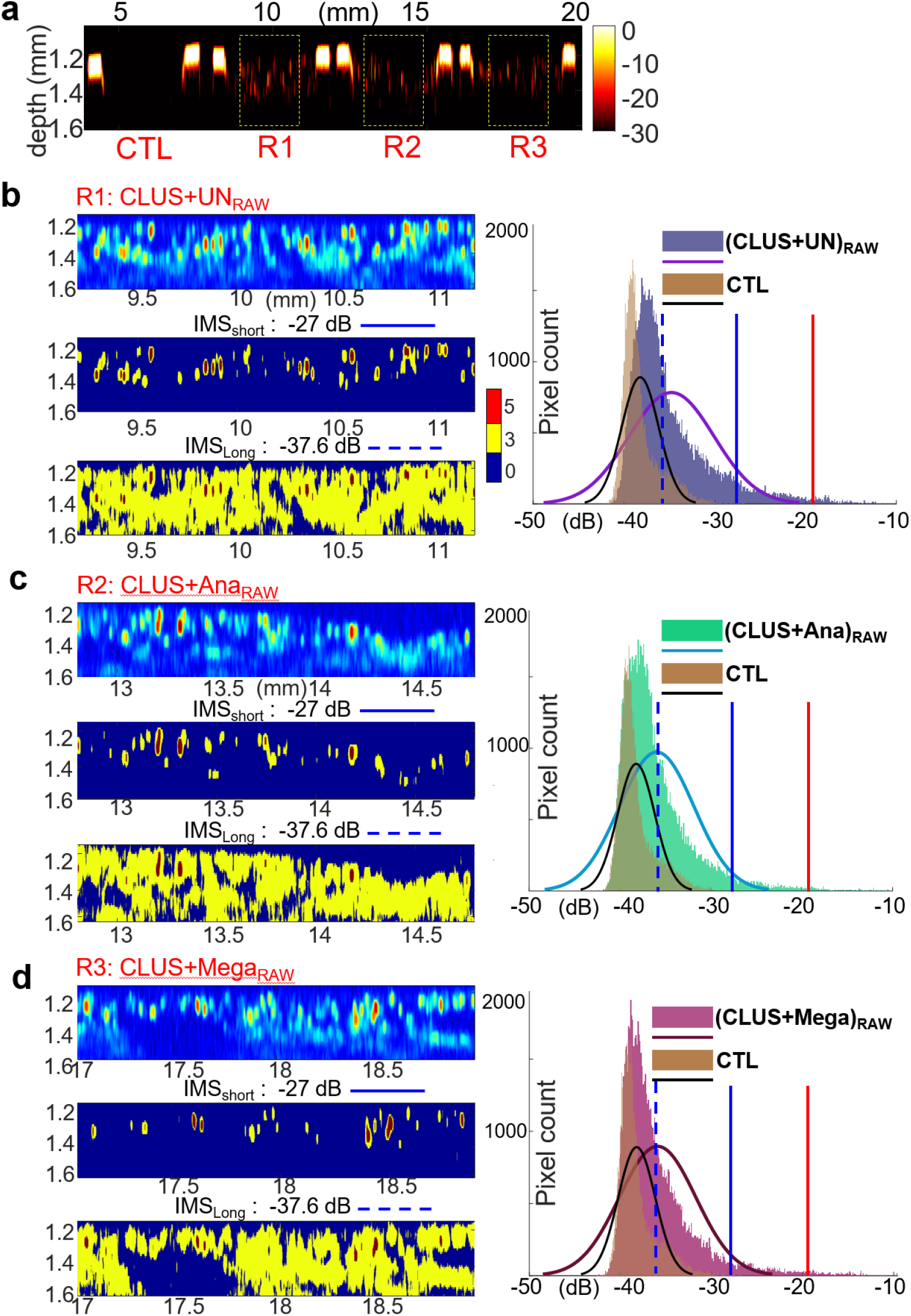
Multiplexed MBF spectral images of mixed RAW cells. **a.** A MBF spectral image of a mixture of CLUS_RAW_ and UN_RAW_ (CLUS+UN_RAW_) at R1, CLUS_RAW_ and Ana_RAW_ (CLUS+Ana_RAW_) at R2, and CLUS_RAW_ and Mega_RAW_ (CLUS+Mega_RAW_) at R3 was obtained. **b-d.** Zoom-in MBF images of yellow boxes in (**a**) are at the first rows. Histogram compares the number of pixels of MBF signals from CTL and CLUS+UN_RAW_, CLUS+Ana_RAW_, CLUS+Mega_RAW_. Red vertical lines indicate CMS value. Blue solid and dashed lines indicate IMS_short_ and IMS_Long_ values calculated from the mean and standard deviations of MBF signals of R1, R2, and R3. Second and third rows in (**b-d**) represent multiplexed images with IMS_short_ and IMS_Long_. Same CMS were used. Red regions indicate the locations of CLUS_RAW_ and yellow regions indicate the locations of UN_RAW_, Ana_RAW_, and Mega_RAW_. IMS_Long_ includes more pixels than IMS_short_ does, which indicates that signals from background may indicated as GV signal. This is similar to autofluorescence in fluorescence microscopy. IMSεhort and IMS_Long_ corresponds to exposure time control in fluorescence microscopy.

## Discussion

Direct observation of spatial distribution of cells and interaction between cells in their natural three-dimensional environment in vivo provides fundamental understanding of the roles of specific sub-types of cells during disease progression and prognostic biomarkers for patients. Intravital imaging using confocal microscopy, multiphoton microscopy, and nanoscopy is a powerful technique to resolve fine structures at single cell to nanometer resolution, but confocal microscopy has photocytotoxicity and limited imaging depth less than 100 μm due to autofluorescence and scattering. Multiphoton microscopy has deeper penetration depth than confocal microscopy, but usually the penetration is less than 300 μm, preventing real-time deep tissue imaging. CT / MRI / nuclear medicine imaging do not have a single cell resolution, despite deep tissue penetration. Powerful single cell profiling methods, such as imaging mass cytometry by time-of-flight (cyTOF), advance the study of the mechanisms of tumor microenvironment (TME) in contributing to tumor progression; however, CyTOF requires tissue fixation and lacks real-time tracking ability. Collectively, imaging modalities lack deep penetration and/or real-time high spatial resolution to resolve behavior and the spatial distribution of cells within wound sites and TME. Multiplexed imaging capability in fluorescence microscopy using fluorescent proteins such as GFP and RFP has been widely used to identify multiple organelles within single cells or differentiate different types of cells under consideration. However, the intrinsic challenges in optical imaging has been shallow penetration depth.

To address these challenges, we developed a multiplexed ultrasound imaging approach using clustered and unclustered GVs as ultrasound contrast agents and MBF spectral imaging with pseudo-coloring scheme. Clustered GVs and individualized and unclustered GVs were isolated from their host bacteria. To isolate AnaGVs, usually it takes four weeks and SGVs isolation takes two weeks with no cytotoxicity. We demonstrate that the growth temperature and duration were important for GV formation. We chose urea to separate clustered SGVs due to urea’s stable performance. We demonstrated that MBF spectral imaging specifically visualizes the location of clustered and unclustered GVs using 130 MHz and 50 MHz ultrasound when GVs are phagocytosed in murine macrophages. CLUS and unclustered GVs such as UN, Ana, and Mega exhibit distinct echo spectrum that can be clearly differentiated as a multiplexing source. This multiplexed MBF spectral imaging approach implemented with clustered and unclustered GVs offers an innovative strategy to acquire the spatial and temporal information about the locations of different species of cells. We demonstrate the application of this approach in visualizing vessels in murine liver specimens and spatial distribution of cells with different GVs as multiplexing source, proving utility in identifying cell locations and small vessel structure. To maximize the multiplexed imaging potential, single cell was used as imaging unit instead of each GV. Here, single cells are containing large amount of GVs in their cytoplasm. After cells phagocytose GVs, GV’s shape and sizes were preserved to provide evident multiplexed imaging capability. Single cell imaging instead of individual GV imaging has three advantages. The size of targets becomes comparable to the wavelength of 130 MHz and 50 MHz ultrasound, resulting in distinct and appropriate MBF spectral signals. GVs expressed or internalized in single cells can be used as beacons and carriers for this multiplexed and single cell imaging by avoiding reticuloendothelial system 21 as cells are not phagocytosed in Kepffer cells in liver. Only small amount of GVs are required because GVs are concentrated in cells to emit MBF spectral signal for imaging. Moreover, long-term tracking of target cells may be performed because of the stability of GVs as we demonstrated that GVs are intact after the overnight incubation in 37°C with 5% CO_2_. This approach may be compatible with live cell imaging in vivo at depth real time using commercially available ultrasound system and drug delivery method using GVs containing therapy drugs.

GV-based MBF spectral imaging with pseudo-coloring scheme visualizes three-dimensional structure in tissue specimen and efficiently rejects signals from out of focus area that resembles confocal microscopy with improved axial resolution with sectioning capabilities. Different filter setups such as IMS_short_ and IMS_Long_ represents different patterns of the locations of unclustered GVs which is similar to longer exposure time in fluorescence microscopy.

In summary, we envision that different species of GVs may be further developed as therapeutic and diagnostic contrast agents due to their stability in circulation system in animals and cytoplasm of incubated cells. This article studies several types of GVs to apply them as ultrasound contrast agents. This could have great implications to understand the bacterial GVs system and to investigate novel microbiome studies.

## Methods

### Gas vesicle preparation

Three different types of GVs were generated and used in this study. We cultured and isolated AnaGVs, MegaGVs, and SGVs from *Anabaena flos-aquae, E. coli* using pST39 plasmid containing pNL29, and *Serratia*^21,28,32^. Bacterial strains and plasmids used in this study are listed in Supplementary Table 1.

### SerratiaGVs

We optimized Serratia culture and SerratiaGV isolation protocols (Fig. 2c and Supplementary Materials and Methods). *Serratia* sp. strain ATCC 39006 were grown in Nutrient Broth medium. 5 ul of the glycerol stock was transferred to 5 ml of culture medium and incubated at 26 °C and 150 rpm for 24 hours. Inoculate 0.1 ml of the orange or yellow suspension into 50 ml tube and grow bacteria in the incubator at 26 °C, 150 rpm for 2-3 weeks dark cycle. Transfer 20 ml culture medium to 5 ml tubes and centrifuge at 350 g at 8 °C for 4 hours. To lyse the Serratia bacteria, add 2 ml of SoluLyse-Sodium Phosphate reagent (Genlantis) per 2 ml of pooled culture, 400 μg/ml lysozyme, 1 μl/ml DNase I (200 Unit/μl) and 100 μl/ml B-PER reagent. Gently rotate the tube for 30 min at room temperature. Centrifuge the tube again at 350 g and 8 °C for 4 hours. Resuspend the layer with 1x PBS and resuspend by centrifugation. This cleaning step needs to be repeated at least 3 times. Harvested GVs were used for experiments in clustered and unclustered states. 1ml of OD_500_= 20 SerratiaGVs can be isolated from 150 ml of culture.

### Unclustering process

Serratia GVs is clustered after isolation. To uncluster them, GV containing solution is mixed with urea at 6 M final concentration, and the resulting solution is gently rotated for 1 hours at room temperature. GV solution with 6 M urea was stored at 4 degrees for 3 days. Resuspend the GV solution with the 1x PBS and centrifuge at 350 g 8 °C for 4 hours. Repeat this process 2 times. Unclustered SerratiaGV concentration decreased by 1/3. Unclustered SerratiaGVs became clustered 3 days after unclustering.

### Cell culture and GV internalization

RAW 264.7 (ATCC) cells were cultured in DMEM supplemented with 10% fetal bovine serum (FBS) and 1% penicillin/streptomycin (P/S) at 37 °C in a humidified atmosphere with 5% CO_2_ and passaged at or before 70% confluence.

To internalize UN, CLUS, Ana, and Mega GVs in RAW cells, cells were cultured in Culture-Inserts 2-well (ibidi) in 35 mm dish 24 hours before incubation of RAW cells with GVs. GVs at OD_500_ of 1 was incubated with RAW cells for 2 hours by inverting the plate for maximum contact between cells and GVs. After incubation for 2 hours, trypsinized cells and washed cells with PBS. A phantom with RAW cells with UN, CLUS, Ana, and Mega was prepared for ultrasound imaging. PCI was taken using an Olympus CX43 microscope with a 100x oil-immersion lens, with a U-TV1XC camera, and LC micro software.

### XTT assay

This analysis is to investigate the effect of prodigiosin and carbapenem produced by Serratia on apoptosis. After inoculating 0.1 ml of the orange or yellow suspension into 50 ml tube, SerratiaGVs were isolated three days, one week, two weeks, and three weeks after the incubation at 26 °C, 150 rpm. The cytotoxicity of GVs towards RAW 264.7 cells were assessed using the XTT (Roche) cell proliferation assay. Cells were seeded into 96-well plates at a density of 2000 cells per well. After RAW cells were culture for 24 hours, RAW cells and GVs were mixed in each well at the final GV concentration OD_500_= 1. Prepare a mixture of 5 ml XTT labeling reagent and 100 ul electron coupling reagent. The mixture was incubated with RAW with GVs at 37 °C for 4 hours. Absorbance was measured using a microplate reader (Tecan) at 492 nm.

### MBF Phantom

Cylindrical fixtures were made with 3.2 mm diameter and 5 mm height for scanning using UBM system as shown in Supplementary Fig. 2C. Prepared GVs were mixed with 1% agarose gel by mixing 0.3 g agarose powder and 30 ml Type I water. Agarose gel solution is pre-warmed at 50°C and taken from the working solution to make GV phantom. GVs of 30 μl and 1% agarose gel of 30 μl were pipetted three times and 38 μl of mixture was taken and immediately placed in the cylindrical fixture. Cylindrical fixtures containing GV phantom were placed at 4°C refrigerator for two minutes before ultrasound scanning. For RAW cell with GV experiments, the same procedure was followed to make RAW cell phantom.

### UBM microscopy

A schematic diagram of ultrasound biomicroscopy (UBM) system for the scanning of MBF phantoms or mouse liver specimen are shown in Supplementary Fig. 2b. A 3D axis stage and a water cuvette were placed on an optical board. The 3D axis stage precisely controls the location of the high frequency ultrasonic transducer (grey color) to perform raster scan or 1D scan with the step size of 1 μm. 130 MHz or 50 MHz transducer was connected to a pulser / receiver and an analog-to-digital (A/D) card for radiofrequency (RF) data acquisition. LabView custom program controls the sequence between pulse / receiver and stage control to acquire A-mode signal for offline post-processing. Custom built Matlab program was used to calculate MBF spectral signals and reconstruct MBF 2D images and B-mode images. Along the width direction (Supplementary Fig. 2c), 20 mm was scanned with 1 μm step size. For mouse liver specimen scanning, elevation (z) direction scanning was also performed with a step size of 5 μm (Supplementary Fig. 4).

### Ultrasonic transducer development

Lithium niobate (LiNbO_3_, LNB, PZT) was used to develop 130 MHz and 50 MHz transducers as shown in Supplementary Fig. 2a. Detailed protocols are available in our papers. Briefly, two 36-deg-rotated Y-cut LNB were lapped down to 20 μm and 50 μm for 130 MHz and 50 MHz ultrasonic transducers, respectively. A conductive silver epoxy was cast on to the lapped LNB as a backing material. Each piece of LNB with back material was turned down to 1 mm and 3 mm in diameters for 130 MHz and 50 MHz transducers, respectively. After these cylindrical shaped LNB stack was placed in bronze tubing and fixed with insulating epoxy permanently. Using spheres with a dimeter of 2 mm and 7 mm, the apertures of 130 MHz and 50 MHz were press-focused to have *f_number_* of 1.5 and 1.16, respectively. After connecting the backing and the positive wire of SMA connector, 2 μm of parylene coating was deposited on whole aperture as a matching layer. Using UBM system, time domain pulse and echo signal was obtained and their frequency domain response was calculated (Supplementary Fig. 2a). Center frequency and −6 dB bandwidth were calculated.

### Transmission Electron Microscopy (TEM)

#### GVs TEM imaging

TEM micrographs of GVs were obtained using a JEOL TEM 2011, fitted with the Gatan Digital Micrograph software. An aliquot (5 μl) of each sample was placed onto a CF300-Ni mesh Nickel grid (Electron Microscopy Sciences, EMS). The sample grid was deposited for 1 minute using 2% uranyl acetate solution, then stains were removed with filter paper, and dried for 24 hours before TEM image^19,32^. The samples were imaged using a low electron dose rate of 120 kv to avoid sample damage. All the registered TEM micrographs were analyzed in ImageJ.

#### Cell TEM imaging

Cells were grown and washed as in above (Cell culture and GV internalization). After GVs were phagocytosed by RAW cells, trypsinized cells were resuspended in 4% paraformaldehyde and washed with 1x PBS before imaging. Fixed RAW cells were directly imaged without staining. A JEOL TEM 2011 TEM with an accelerating voltage of 120 kV was used for both low- and high-resolution TEM measurements. All measurements were performed on the lacey holey TEM grids from Carbon film (EMS).

#### Confocal microscopy imaging

A mixture of 1-1-(3-di-methylaminopropyl)-3-ethylcarbodiimide hydrochloride (EDC) (2mM) prepared in PBS and sulfo-N-hydroxysuccinimide (sulfo-NHS) (5 mM) dissolved in a volume of 5 μl Alexa Fluor 488 carboxylic acid was rotated for 30 min at room temperature in the dark, in order to activate the carboxyl groups of GVs. The activated Alexa 488 solution (10 μl) was then added to the GV suspension, and the final mixture was rotated overnight in the dark at 4 °C. Fluorescently labeled GVs were purified using dialysis against PBS36. Alexa 488 conjugated GVs were centrifuged at 4 °C, 300 g. The concentration of Alexa 488 labeled GVs was adjusted at OD_500_= 1 (OD_500_=20, 50 ul/PBS 950 ul).

Since GVs are floating in liquid, RAW cells and GVs labeled with Alexa 488 were incubated in Culture-Inserts 2-well (ibidi) attached on a 24-well plate at 37 °C for 2 hours. The cells were washed twice with PBS and then stained with 50 μl of CellMark Orange plasma membrane stain (Thermo Fisher Scientific) in the dark at room temperature for 15 min. The cells were washed three times with 100 μl of 1× PBS followed by trypsinization and washing with PBS. Cells were fixed with 4% paraformaldehyde for 20 min and permeabilized with 0.5% Triton X-100 for 15 min. Cells were counterstained in 1 μg/ml DAPI and mounted in mounting medium. Images were obtained using a confocal microscope (Nikon A1R-MP Laser Scanning Confocal Microscope) and processed using Nikon analysis software (Nikon). Confocal z-stacks of 0.3 μm were acquired and images were analyzed with Nikon analysis software.

#### Mouse liver tissue preparation for imaging and histology

All imaging experiments were performed on 8-week-old female mice using protocols approved by the Institutional Animal Science and Use Committee of the University of Notre Dame. Mice were fed a standard chow diet and were kept on a 12-hour light / 12-hour dark cycle at 22 ± 2°C with free access to food and water. Mice were sacrificed after anesthesia with Avertin (Tribromoethanol), and intrahepatic perfusion was performed with 50 ml saline. After perfusion, 1 ml of OD_500_= 20 AnaGV and SGV were injected into two mice, separately. Livers with AnaGVs and SGVs were immediately isolated and immersed into 10% formalin for 24 hours for tissue fixation. To image mouse livers injected with AnaGVs and SGVs, tissue specimens were placed on a gelatin phantom cross-linked with paraformaldehyde after degassing and cooling.

#### Histology

The hematoxylin and eosin (H&E) staining and immunohistochemical analysis were performed using paraffin sections, as described previously37. Briefly, Mouse liver tissue was fixed overnight in 10% formalin, dehydrated, paraffin incubated, and sectioned at 5 μm. A standard H&E protocol was used for staining afterwards (95% EtOH, 70% EtOH, H2O for 30 s each, hematoxylin for 3 min, H2O, 70% EtOH, 95% EtOH for 30 s, eosin for 1 min, 95 and 100% EtOH for 30 s each, xylene for 2 min). Images were obtained with Nikon AZ 100 Macro/Zoom Microscope.

#### Statistics

Statistical analyses were performed using Matlab and GraphPad Prism 5 software (GraphPad Software, La Jolla, CA, USA).

## Acknowledgements

We thank S. Chapman and S. Cole for their technical assistance with histology experiments. M. Zhukovskyi for help with TEM imaging. We appreciate the help of Q. Wang with animal protocols. This work was supported in part by National Institute of Health (NIH) grant No. GM120493 and National Science Foundation (NSF) grant No. CBET 1943852, and Harper Cancer Research Institute Cure Cancer Venture (CCV) grant.

## Author contributions

S.Y. and S.K. conceived the designed the study. S.K. grows bacteria, isolated GVs, performed TEM, confocal, PCI imaging, histology, grow cells and incubated cells with GV. S.Y. developed the system and ultrasonic transducers, performed phantom and mouse imaging using UBM system and analyzed data and constructed images. S.Z. provided and consulted mouse models. S.K and S.Y. analyzed the results. S.Y. and S.K. wrote the manuscript. All authors reviewed and agreed the content of the manuscript.

