## Supplementary information for "Multiplexed ultrasound imaging using clustered gas vesicles and spectral imaging"

### Materials and Methods

**Supplementary Table S1.** List of bacterial strains and plasmids used in this study.

| Strain or plasmid | Genotype or description | Source |
|---|---|---|
| *Serratia* sp.[1] | Cat. no. 39006 | ATCC |
| *Anabaena flos-aquae*[2] | Cat. no. CCAP 1403/13F | Culture Collection of Algae and Protozoa (CCAP) |
| Rosetta 2(DE3) pLysS competent *E. coli*[2] | Cat. no.71401-4 | Sigma |
| pST39 plasmid containing pNL29 Mega GV gene cluster[2,3] | ID no. 91696 | Addgene |

#### AnaGVs

AnaGVs was isolated as described previously[2,3]. Briefly, Ana was incubated in Gorham medium containing BG-11 solution in a 2 L flask for 14 hours photoperiod and 10 hours dark at 25 °C, 100 rpm shaking. After 3 weeks, the culture was transferred to a separatory funnel, and the buoyant culture was separated after 48 hours. To isolate AnaGV, a total of 500 mM sorbitol and 10% Solulyse (Genlantis) solution was added and incubated for 8 hours. The suspended layer was resuspended in 1X phosphate buffered saline (PBS) by centrifugation. After washing repeatedly, GV, which is a white layer, was acquired. GV concentrations were determined by measuring OD at 500 nm ($OD_{500}$) using NanoDrop 2000 UV-Vis Spectrophotometer (Thermo Fisher Scientific).

#### MegaGVs

MegaGVs were expressed in *E. coli* Rosetta 2 (DE3) pLysS cells (Novagen) and purified as described previously[2,3]. MegaGV was generated by heterologous expression in *E. coli*. pST39 plasmid containing pNL29 were transfected into Rosetta. Transformed *E. coli* was contained in LB culture medium with 1x ampicillin, 1x chloramphenicol, and 1% glucose. When the $OD_{600}$ value reached 0.4-0.6, IPTG was added and cultured for 24 hours. To isolated Mega GV, growth media was centrifuged at 500g and 25°C in 50 ml conical tube. Transparent portion of the centrifuged medium was carefully removed. MegaGVs were in the pellet and supernatant except for the intermediated layer. To separate GVs from cells in the pellet and supernatant, after adding SoluLyse-Tris solution, lysozyme, and DNaseI to 50 ml conical tube with GVs, incubate at room temperature for 10 minutes. Centrifuge lysate including GVs sample in 50 ml at 350g and 8°C for 4hours. Resuspend the white layer with 1x PBS and resuspend by centrifugation. After repeating this process at least 3 times. By applying 6M urea, uncluster MegaGVs.

#### Serratia bactseria growth

Serratia growth under various conditions (Fig. 2c) was measured at OD at 600 nm with the NanoDrop 2000 UV-Vis Spectrophotometer. Bacteria were inoculated with growth media at 1:500 dilution form the pre-culture. Three flasks were grown at 20°C, 26°C, and 30°C incubation condition. Bacterial optical density ($OD_{600}$) was measured every 1 hour to observe growth curve. Phase contrast imaging (PCI) with 100x magnification was used to image Serratia proliferation and GV yield. Serratia were imaged with Olympus CX43 microscope using with a U-TV1XC camera and LC micro software. TEM imaging was performed to observe Serratia and GV formation at 13 hours, 39 hours, and 72 hours after the incubation started. Length and width of GVs were measured using ImageJ software.

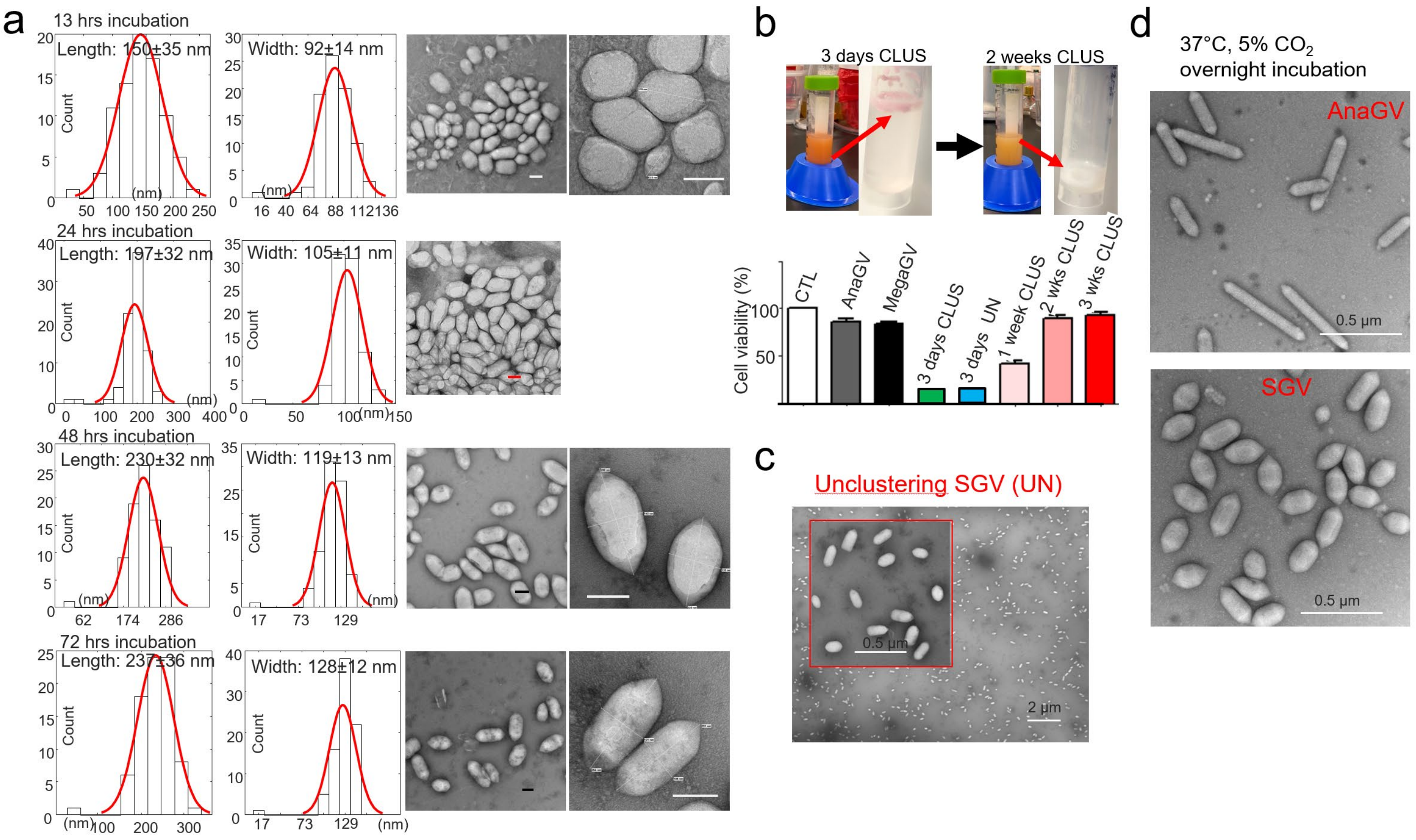


**Supplementary Figure 1. a**. Gas vesicles (GV) grow in Serratia and the maximum sizes reach 48 hours after the incubation at 26°C. TEM images show Serratia GVs (SGV) at different incubation time. Scale bars indicate 100 nm. **b**. Serratia produces prodigiosin and carbapenem[1,4]. Prodigiosin is red pigment. Long term incubation of Serratia more than 2 weeks removes prodigiosin and carbapenem. Isolated GV clouds changed from red to white. XTT assay indicates that GVs from Serratia after long term incubation did not influence RAW cell viability. **c**. After urea treatment, SGVs were unclustered. TEM image of unclustered SGVs. **d**. GVs were stable and stayed intact after overnight incubation at 37°C.

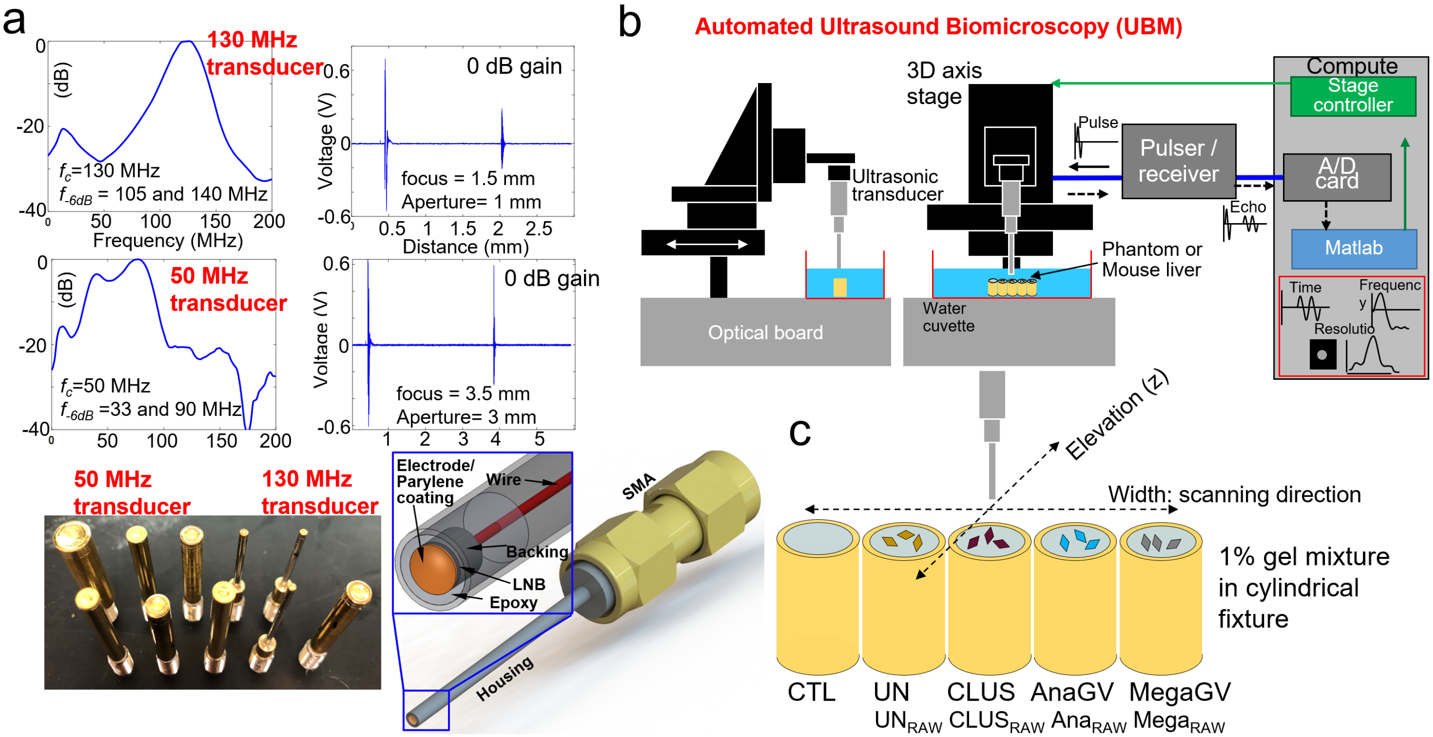


**Supplementary Figure 2. a**. Echo spectrum and time responses of in-house developed transducers present that the center frequencies were 130 MHz and 50 MHz. Lithium niobate (LNB) was lapped down to desired thickness and electrode and parylene coating were applied. Acoustic stack was fixed with epoxy at the tip of housing. SMA connector was used for electrical connection. **b**. Ultrasound biomicroscopy (UBM) consists of 3D translation stages, pulser / receiver, A/D card, and stage controller. 3D translation stages control the scanning of a phantom or mouse liver. **c**. 130 MHz or 50 MHz were used to scan GV and RAW cell phantoms along width direction using UBM.

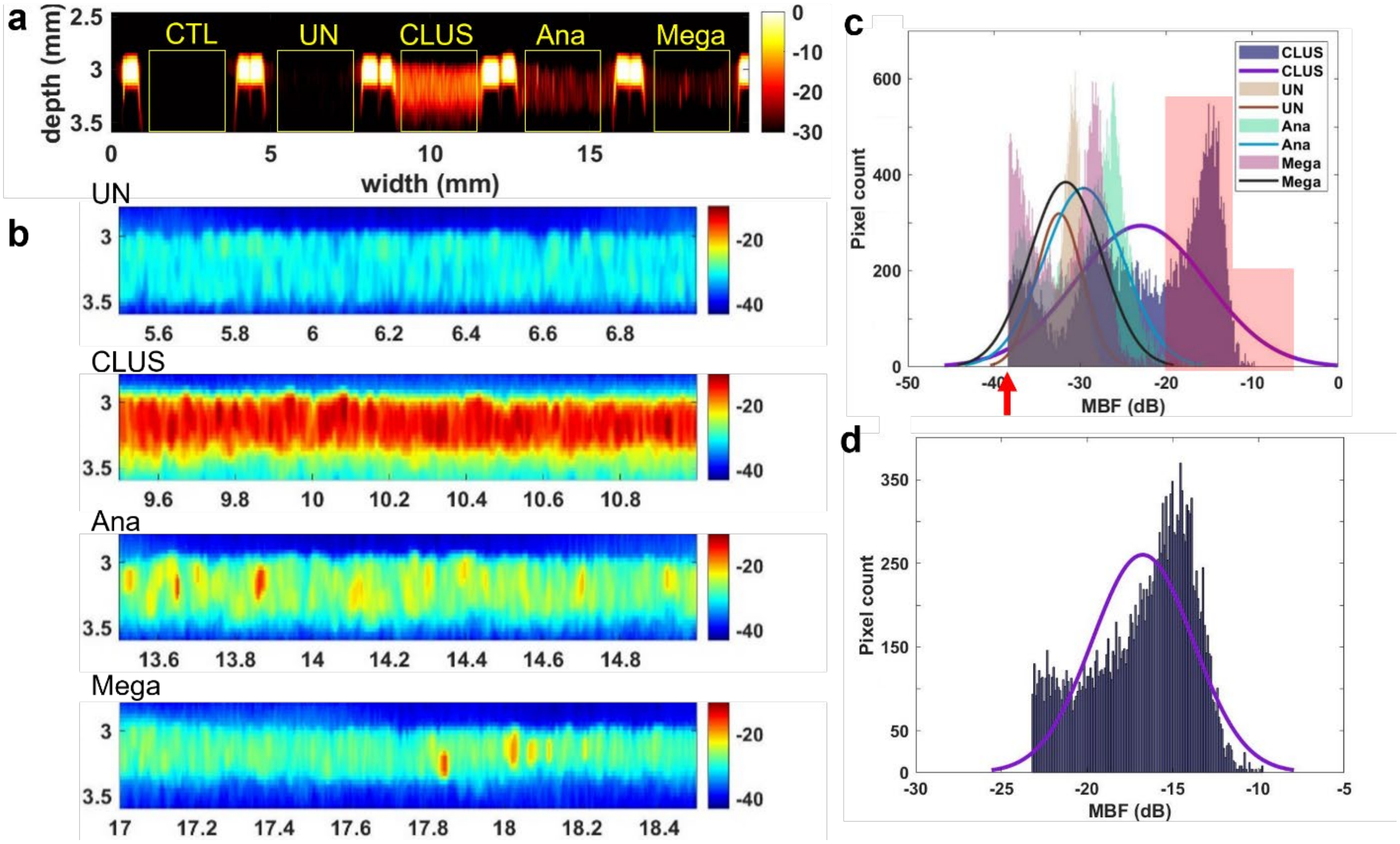


**Supplementary Figure 3. a**. MBF spectral images of control (CTL), UN, CLUS, Ana, Mega GVs scanned by 50 MHz ultrasound. GVs are mixed with 1% gelatin at the final $OD_{500}$ of 10. **e.** Zoom-in MBF spectral images of yellow boxes in **a** indicates that CLUS signal is higher than other MBF from other GVs. **c.** Histogram of MBF signal of all pixels in yellow boxes shows that CLUS signal has distinct MBF signal indicated by red region. Before counting MBF signals of each pixel, all pixels with MBF signals lower than basal level was removed (red arrow) **d.** Histogram of MBF signal of CLUS distinct from UN, Ana, and Mega is shown. This histogram was generated after removing the mean of MBF signals from UN, Ana, and Mega. At this $OD_{500}$ of 10 imaged by 50 MHz ultrasound, the mean and the standard deviation of MBF signal of CLUS is -16.8 ± 3.0 dB.

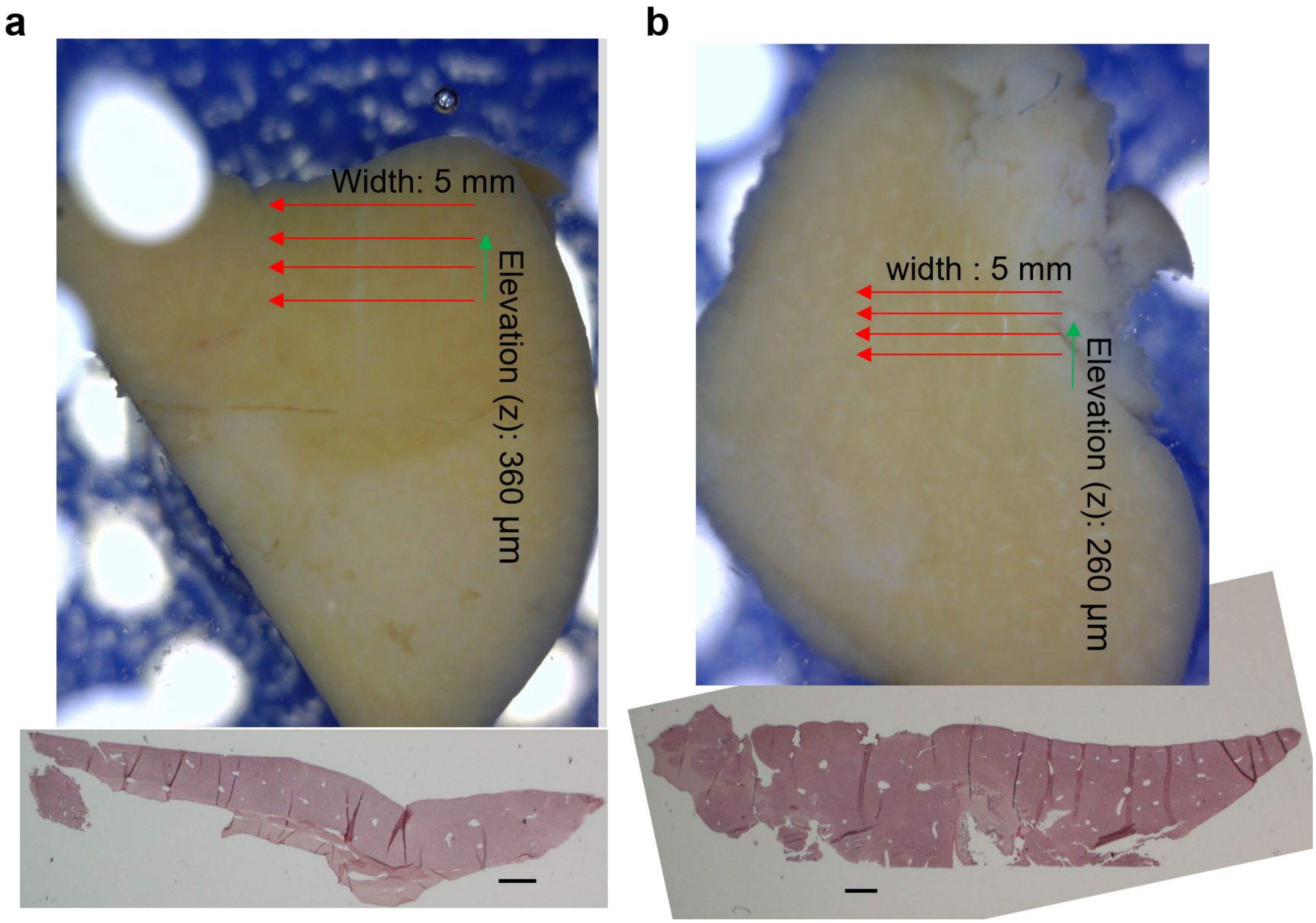


**Supplementary Figure 4.** Mouse liver was injected with **a** AnaGVs and **b** SGVs. Tissue images and the hematoxylin-eosin (H&E) staining in paraffin sections are shown. H&E images show hollow regions which indicate vessels in liver. 2D section images were obtained by scanning 5 mm along the width direction. 73 and 53 sections were imaged along elevation direction with 5 μm intervals to cover 360 μm and 260 μm regions for **a** AnaGVs injected and **a** SGV injected mouse livers, respectively. 3D rendering was produced shown in Fig. 5. Scale bars:1 mm.

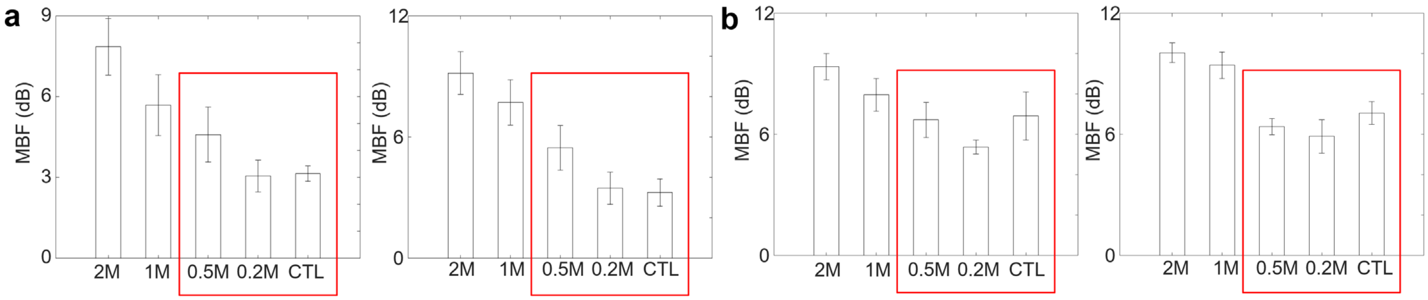


**Supplementary Figure 5.** MBF signal of RAW cells without GVs measured by **a** 130 MHz and **b** 50 MHz ultrasound. 0.2 million (M), 0.5 M, 1 M, and 2 M cells were used.

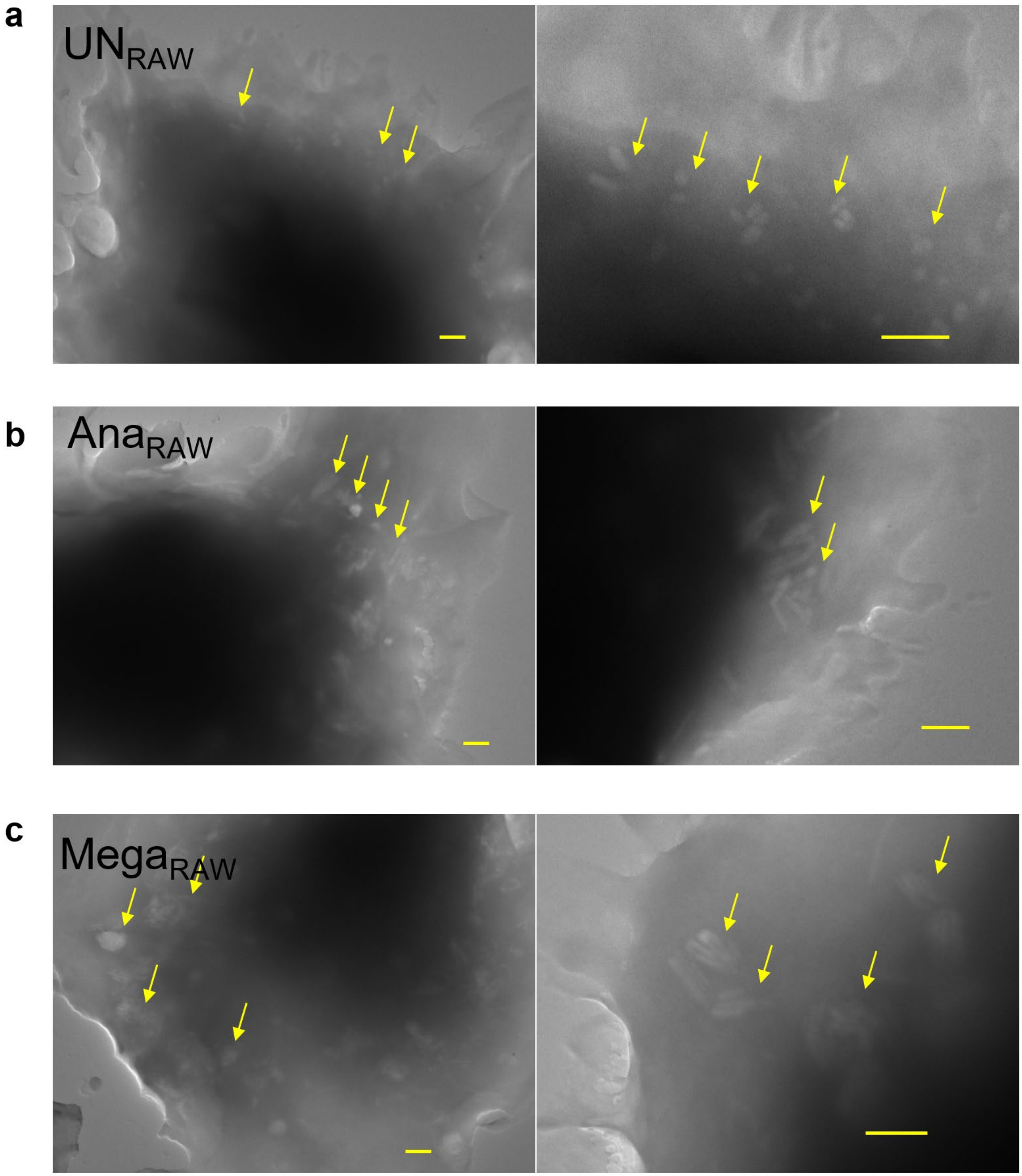


**Supplementary Figure 6.** TEM images confirm non-clustered GV internalization in RAW cells (yellow arrows) after two hours of incubation at 37°C. **a** Unclustered SerratiaGVs, **b** AnaGVs, and **c** MegaGVs were used. Scale bars: 500 nm.

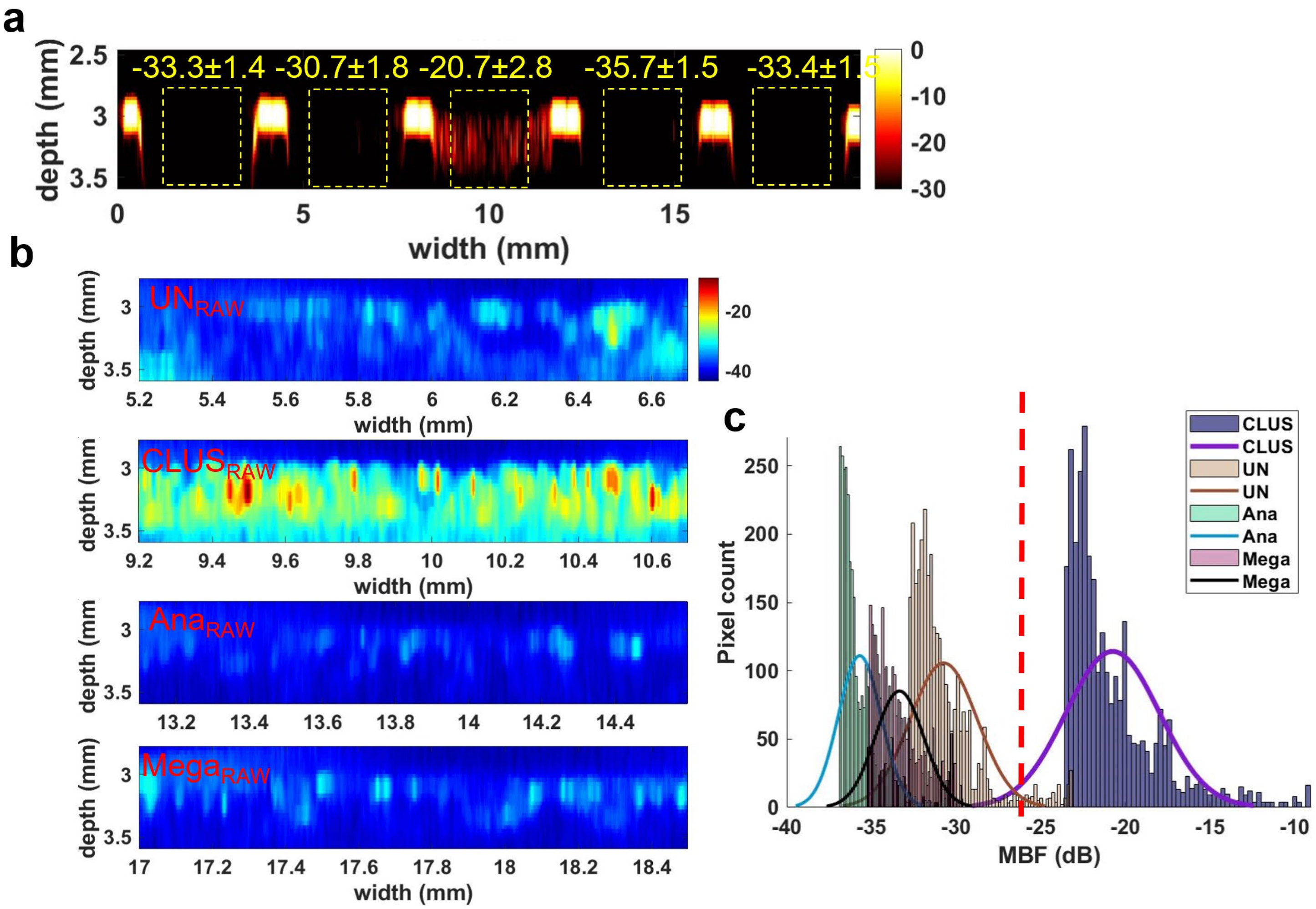


**Supplementary Figure 7. a.** MBF spectral images of RAW cells (CTL) and RAW cells with UN ($UN_{RAW}$), CLUS ($CLUS_{RAW}$), Ana ($Ana_{RAW}$), and Mega ($Mega_{RAW}$) using 50 MHz ultrasound. Means and standard deviations of MBF signal from $CLUS_{RAW}$ are distinct from $UN_{RAW}$, $Ana_{RAW}$, $Mega_{RAW}$., which shows distinct pixel counts in **c** the histogram. **b.** Zoom-in MBF spectral images of $UN_{RAW}$, $CLUS_{RAW}$, $Ana_{RAW}$, and $Mega_{RAW}$ indicates the distribution of GVs internalized by RAW cells. Error bars indicate ± one standard deviation.

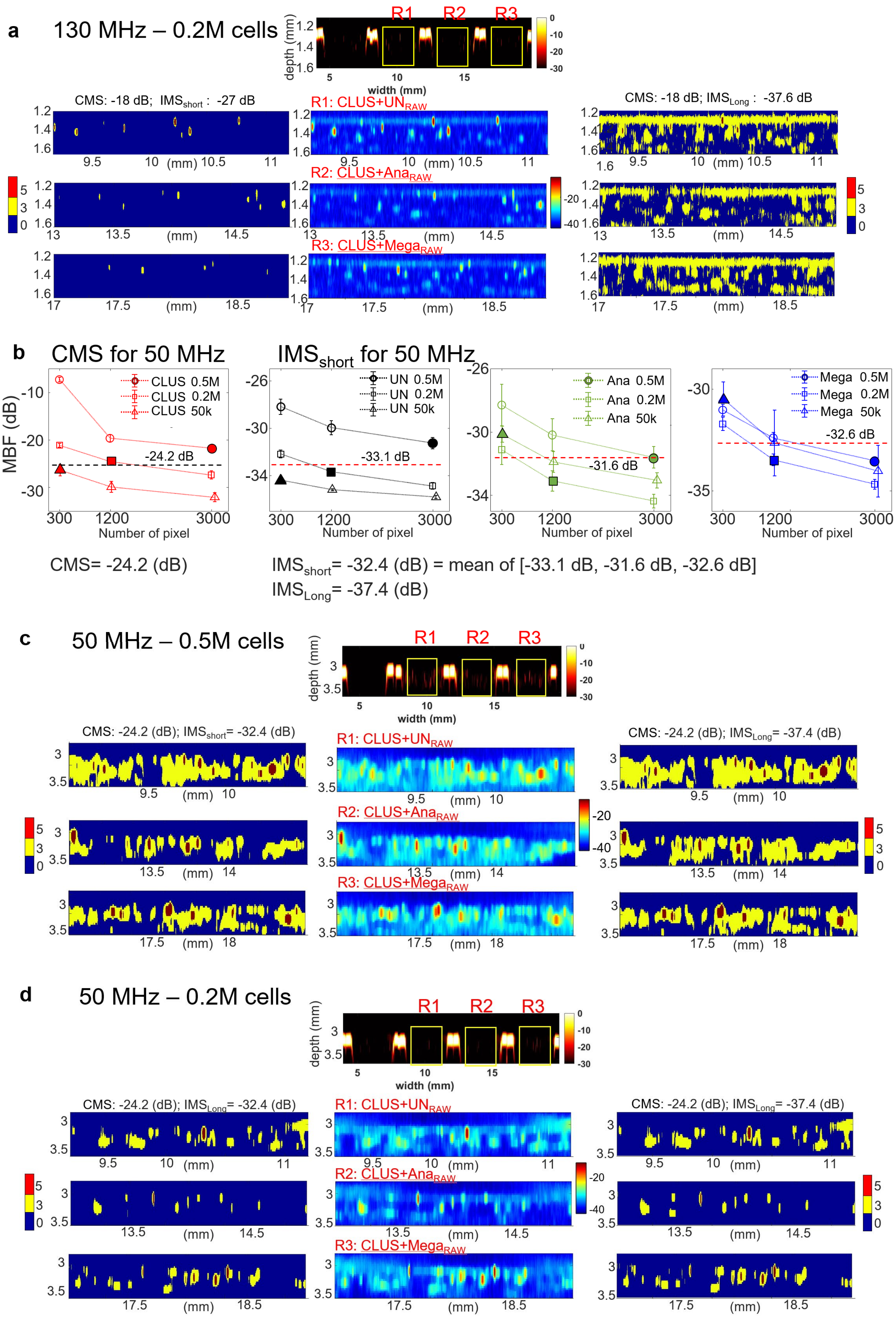

a
130 MHz – 0.2M cells
R1
R2
R3
depth (mm)
width (mm)
CMS: -18 dB; IMSshort : -27 dB
R1: CLUS+UNRAW
CMS: -18 dB; IMSLong : -37.6 dB
R2: CLUS+AnaRAW
R3: CLUS+MegaRAW
b
CMS for 50 MHz
IMSshort for 50 MHz
MBF (dB)
CLUS 0.5M
CLUS 0.2M
CLUS 50k
-24.2 dB
UN 0.5M
UN 0.2M
UN 50k
-33.1 dB
Ana 0.5M
Ana 0.2M
Ana 50k
-31.6 dB
Mega 0.5M
Mega 0.2M
Mega 50k
-32.6 dB
Number of pixel
CMS= -24.2 (dB)
IMSshort= -32.4 (dB) = mean of [-33.1 dB, -31.6 dB, -32.6 dB]
IMSLong= -37.4 (dB)
c
50 MHz – 0.5M cells
CMS: -24.2 (dB); IMSshort= -32.4 (dB)
CMS: -24.2 (dB); IMSLong= -37.4 (dB)
d
50 MHz – 0.2M cells
CMS: -24.2 (dB); IMSLong= -32.4 (dB)
CMS: -24.2 (dB); IMSLong= -37.4 (dB)

**Supplementary Figure 8. a.** Mixtures of $CLUS_{RAW}$ and $UN_{RAW}$ (CLUS+$UN_{RAW}$) at R1, $CLUS_{RAW}$ and $Ana_{RAW}$ (CLUS+$Ana_{RAW}$) at R2, and $CLUS_{RAW}$ and $Mega_{RAW}$ (CLUS+$Mega_{RAW}$) at R3 were scanned by 130 MHz ultrasound with 0.2M cells. Zoom-in MBF images of R1, R2, and R3 regions are in the second column. The first and the third columns represent multiplexed images with $IMS_{short}$ and $IMS_{Long}$. Same CMS were used. **b.** CMS for 50 MHz ultrasound multiplexed imaging was defined by taking the mean of MBF signal from 0.5M, 0.2M, 50000 $CLUS_{RAW}$ using 3000, 1200, and 300 pixels (solid circle, square, and triangle) by considering the number of cells within the imaging slice. Similarly, $IMS_{short}$ was defined by the mean of means of MBF signals (arrow head and red dashed lines) from $UN_{RAW}$, $Ana_{RAW}$, and $Mega_{RAW}$. Error bars indicate ± one standard deviation. **c-d.** Mixtures of $CLUS_{RAW}$ and $UN_{RAW}$ (CLUS+$UN_{RAW}$) at R1, $CLUS_{RAW}$ and $Ana_{RAW}$ (CLUS+$Ana_{RAW}$) at R2, and $CLUS_{RAW}$ and $Mega_{RAW}$ (CLUS+$Mega_{RAW}$) at R3 were scanned by 50 MHz ultrasound with **(c)** 0.5M and **(d)** 0.2M cells. Zoom-in MBF images of R1, R2, and R3 regions are in the second column. The first and the third columns represent multiplexed images with $IMS_{short}$ and $IMS_{Long}$. Same CMS were used.
